# KDM6 Enzymes are the Mechanistic Targets of Mutant IDH that Dictate Replication Stress Sensitivity

**DOI:** 10.64898/2026.04.14.715350

**Authors:** Alexander C.-Y. Tsai, Mathew D. Lin, Vinesh T. Puliyappadamba, Dorothy M. Junginger, Victoria G. Donovan, Eleanor G. Kaplan, Laura M. Drepanos, Hiroaki Wakimoto, Daniel P. Cahill, Julie A. Losman, Kent W. Mouw, Kalil G. Abdullah, John G. Doench, Duane Nash, Daniel Vitt, Christian Gege, Hella Kohlhof, Samuel K. McBrayer, William G. Kaelin, Diana D. Shi

## Abstract

Cancer-associated isocitrate dehydrogenase (IDH) mutations sensitize gliomas to replication stress, although the underlying mechanisms are unclear. IDH-mutant enzymes synthesize (*R*)-2-hydroxyglutarate (R2HG), which broadly inhibits 2-oxoglutarate-dependent enzymes. We performed forward genetic screens targeting all 2-oxoglutarate-dependent enzymes and discovered that KDM6 histone demethylases play a vital role in protecting cells from replication stress. Genetic or R2HG-mediated repression of KDM6 catalytic activity sensitized glioma cells to disparate replication stress-inducing drugs, including Ataxia-telangiectasia and Rad3-related (ATR) and dihydroorotate dehydrogenase (DHODH) inhibitors. This liability is generalizable because KDM6A loss-of-function mutations commonly observed in urothelial carcinomas sensitized bladder cancer cells to DHODH inhibition, thereby phenocopying IDH mutations in glioma. To exploit these oncogene-induced replication stress vulnerabilities, we developed an effective, on-target, and well-tolerated DHODH inhibitor, GLIO-1, that is poised for clinical translation. Collectively, we reveal KDM6 activity as a fundamental determinant of replication stress sensitivity and nominate pan-cancer, mechanism-based biomarkers of ATR and DHODH inhibitor efficacy.

**STATEMENT OF SIGNIFICANCE:** We discovered that the KDM6 enzymes are the mechanistic targets of R2HG that mediate mutant IDH-induced replication stress hypersensitivity. We report a promising new DHODH inhibitor, GLIO-1, and nominate KDM6 and IDH mutations as predictive biomarkers for the antitumor effects of GLIO-1 and other replication stress inducers.

## INTRODUCTION

Gliomas are the most common primary malignant brain tumor in adults, a subset of which have point mutations in genes encoding isocitrate dehydrogenase (IDH1/2) metabolic enzymes. IDH-mutant enzymes, the most common of which is IDH1-R132H in glioma, convert 2-oxoglutarate (2OG) to millimolar levels of the oncometabolite (*R*)-2-hydroxyglutarate (R2HG). R2HG, which is structurally similar to 2OG, competitively inhibits most 2OG-dependent enzymes (including metabolic enzymes, DNA demethylases, and histone demethylases) and activates EglN prolyl hydroxylases (1). Mutant IDH therefore causes widespread metabolic reprogramming, histone hypermethylation, and DNA hypermethylation that are thought to promote glioma initiation.

In addition to driving oncogenesis, mutant IDH also confers collateral vulnerabilities that can be exploited therapeutically. For example, mutant IDH has been linked to altered DNA damage responses induced by various therapies including alkylating agents, poly (ADP-ribose) polymerase (PARP) inhibition, poly(ADP-ribose) glycohydrolase (PARG) inhibition, and/or radiation (2–14). We and others have shown that mutant IDH confers sensitivity to drugs that induce replication stress such as inhibitors of the de novo pyrimidine nucleotide synthesis enzyme dihydroorotate dehydrogenase (DHODH) or Ataxia-telangiectasia and Rad3-related (ATR) inhibitors (5,6,15). However, the mechanisms underlying this replication stress sensitivity remain incompletely understood.

In this study, we sought to identify the molecular mechanism by which IDH mutations sensitize gliomas to replication stress. Because R2HG inhibits many 2OG-dependent enzymes, we hypothesized that one or more of these enzymes natively protect cells from replication stress-induced DNA damage and cell death. To identify such enzymes, we performed unbiased, forward genetic screens targeting all 2OG-dependent enzymes in the presence or absence of replication stress-inducing drugs. We identified KDM6 enzymes as the mechanistic targets of R2HG that mediate response to DHODH inhibitors and ATR inhibitors in glioma. We leverage this mechanistic understanding to demonstrate efficacy of a novel, brain-penetrant DHODH inhibitor and report its utility in both IDH-mutant gliomas as well as KDM6-altered bladder cancers.

## RESULTS

### Unbiased genetic screens show that loss of KDM6 sensitizes glioma cells to replication stress

R2HG competitively inhibits many 2OG-dependent dioxygenases. To assess whether modulation of one or more 2OG-dependent dioxygenases accounts for replication stress hypersensitivity of IDH-mutant tumor cells, we used isogenic IDH-mutant and IDH-wildtype stable cell lines created from endogenously IDH-wildtype human oligodendroglioma (HOG) cells or immortalized human astrocytes (NHAs) (9,15). Each line was engineered to express the mutant IDH1-R132H (R132H) oncoprotein or an empty vector (EV) control. We have shown that IDH-mutant cells in these models accumulate millimolar levels of R2HG and display increased sensitivity to replication stress compared to their EV-expressing controls (9,15).

Ascorbate is a requisite cofactor for 2OG-dependent dioxygenases (including TET and KDM enzymes that demethylate DNA and histones, respectively). Previous work demonstrated that ascorbate 2-phosphate (A2P), an ascorbate analog that is converted to ascorbate intracellularly, could reverse transcriptional consequences of epigenetic alterations caused by R2HG (16). Therefore, we asked if A2P treatment rescued replication stress hypersensitivity induced by mutant IDH. If so, this would imply that inhibition of 2OG-dependent dioxygenase(s) by R2HG accounted for this replication stress vulnerability phenotype.

We focused our studies on anticancer drugs that act nearly exclusively through replication stress induction such as the DHODH inhibitor orludodstat (BAY 2402234) and the ATR inhibitor ceralasertib. As expected, both induced cytotoxicity in a replication-dependent manner, as pretreatment with the CDK4/6 inhibitor palbociclib nearly fully rescued the cytotoxicity induced by both drugs (Fig. S1A). We then used BAY 2402234 and ceralasertib in isogenic cell lines and found that exogenous A2P treatment fully rescued mutant IDH-induced sensitivity to these drugs in HOG cells (Fig. 1A–B). This rescue was not due to growth arrest caused by A2P, as A2P treatment did not alter cell proliferation or the proportions of cells in S phase (Fig. S1B and S1C). Ascorbate can suppress reactive oxygen species (ROS), and ROS can contribute to DNA damage. A2P treatment also did not, however, suppress ROS in our drug treatment experiments (Fig. S1D). Therefore, rescue by A2P was not due to modulation of cell cycle kinetics or ROS.

**Figure 1.**
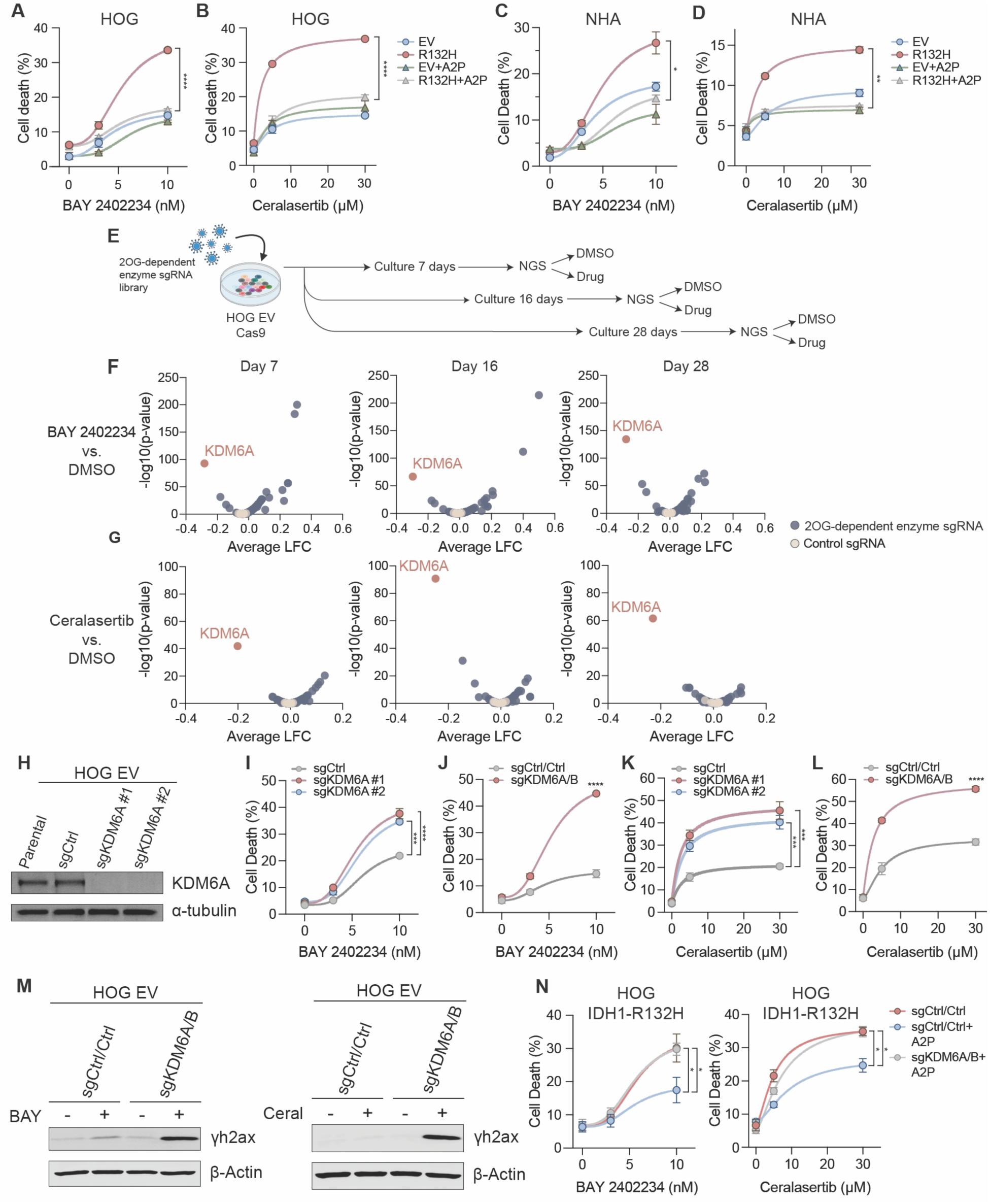
CRISPR screens nominate loss of KDM6 as a sensitizer to replication stress. **(A–B)** Cell death assays of HOG EV or IDH1-R132H (R132H) cells treated with the indicated drugs with or without 200 µM ascorbate 2-phosphate (A2P) (*n =* 3 per treatment condition). (A) 10 nM R132H vs. 10 nM R132H+A2P: p<0.0001; (B) 30 µM R132H vs. 30 µM R132H+A2P: p<0.0001. **(C–D)** Cell death assays of NHA EV or IDH1-R132H cells treated with the indicated drugs with or without 200 µM A2P (*n =* 3 per treatment condition). (C) 10 nM R132H vs. 10 nM R132H+A2P: p=0.03; (D) 30 µM R132H vs. 30 µM R132H+A2P: p=0.0051. **(E)** Schema of CRISPR screens with library of sgRNAs targeting 2-oxoglutarate (2OG)-dependent enzymes. **(F–G)** Volcano plot of log fold changes (LFC) of sgRNAs in 10 nM BAY 2402234 vs. DMSO arms (F) or 30 µM ceralasertib vs. DMSO arms (G) of CRISPR screens depicted in (E) (*n =* 4 per drug). **(H)** Immunoblot analysis of HOG EV cells expressing Cas9 and sgRNAs targeting KDM6A or a control sgRNA. **(I)** Cell death assays of cells in (H) treated with indicated doses of BAY 2402234 (*n =* 3). 10 nM sgKDM6A #1 vs. 10 nM sgCtrl: p=0.0002. 10 nM sgKDM6A #2 vs. 10 nM sgCtrl: p<0.0001. **(J)** Cell death assays of HOG EV cells co-expressing sgRNAs targeting KDM6A and KDM6B or control sgRNAs treated with BAY 2402234 (*n =* 3). 10 nM sgKDM6A/B vs. sgCtrl/Ctrl: p<0.0001. **(K–L)** HOG EV cells in (I–J) treated with ceralasertib (*n =* 3). 30 µM sgKDM6A #1 vs. 30 µM sgCtrl: p=0.0004; 30 µM sgKDM6A #2 vs. 30 µM sgCtrl: p=0.0004; 30 µM sgKDM6A/B vs. 30 µM sgCtrl/Ctrl: p<0.0001. **(M)** Immunoblot analysis of sgKDM6A/B or sgCtrl/Ctrl HOG EV cells treated with BAY 2402234 or ceralasertib. **(N)** Cell death assays of HOG IDH1-R132H cells expressing sgKDM6A/B or sgCtrl/Ctrl with or without 200 µM A2P treated with the indicated drugs (*n =* 4 for BAY 2402234 and *n =* 3 for ceralasertib). 10 nM BAY 2402234: sgCtrl/Ctrl vs. sgCtrl/Ctrl+A2P: p=0.048, sgKDM6A/B+A2P vs. sgCtrl/Ctrl+A2P: p=0.027, sgKDM6A/B+A2P vs. sgCtrl/Ctrl: p=0.71; 30 µM ceralasertib: sgCtrl/Ctrl vs. sgCtrl/Ctrl+A2P: p=0.013, sgKDM6A/B+A2P vs. sgCtrl/Ctrl+A2P: p=0.0172, sgKDM6A/B+A2P vs. sgCtrl/Ctrl: p=0.99. For all panels, data are means ± SEM. For (F) and (G), p-values were determined from an empirically derived null distribution created through bootstrapping of negative control sgRNAs. For all other panels, two-tailed p-values were determined by unpaired t-test.

Similar findings were obtained with IDH-mutant and wildtype NHA cells. In this setting, rescue was observed with long-term (Fig. 1C–D), but not short-term (Fig. S1E), A2P treatment. These kinetics mirror the effect of sensitization by mutant IDH itself in NHAs, where late passage (but not early passage) IDH-mutant cells are sensitized to DHODH inhibition (15). These slow kinetics in NHAs parallel those of epigenetic changed caused by modulation of the activity of chromatin-modifying enzymes (17). These findings support a model in which A2P rescues mutant IDH-driven sensitivity to replication stress by reversing R2HG-dependent inhibition of one or more 2OG-dependent dioxygenase chromatin-modifying enzymes.

To ask whether a specific 2OG-dependent enzyme downstream of mutant IDH modulates sensitivity to replication stress, we employed a CRISPR knockout screen using a focused sgRNA library. We infected Cas9-expressing IDH-wildtype (i.e., EV) HOG cells with a custom sgRNA library targeting 126 genes encoding 2OG-dependent enzymes, including dehydrogenases, dioxygenases, and transaminases (Table S1). To account for possible slow kinetics of epigenetic alterations following gene knockout, we cultured cells for 7, 16, and 28 days before treating with drug (BAY 2402234 or ceralasertib) or DMSO control and subsequently determining sgRNA abundance in each treatment group (Fig. 1E). These two drugs induce replication stress through independent, non-overlapping mechanisms. We reasoned that sgRNAs depleted in the presence of these drugs relative to DMSO would nominate genes (hereafter called “hits”) whose loss sensitizes to replication stress.

In both BAY 2402234 and ceralasertib screens, KDM6A was the top hit at every time point based on sgRNA depletion (Fig. 1F–G, Tables S2–3). KDM6 enzymes (KDM6A, B, and C) are Jumonji C (JmjC) domain-containing histone demethylases that demethylate H3K27me3 and are potently inhibited by R2HG, with IC_50_ values more than ten times lower than TET2 (18), which is a pathogenic target of R2HG in IDH-mutant leukemia (19). However, a functional role of KDM6 enzymes in mediating response to replication stress in glioma has not been described. Importantly, fitness defects conferred by KDM6A loss were specific to drug treatment, as KDM6A sgRNAs were not depleted in untreated cells (Fig. S1F). This latter observation is consistent with KDM6A’s role as a tumor suppressor in cancers including bladder cancer and leukemia (20–23).

Notably, some sgRNAs were *enriched* in drug conditions compared to DMSO control, implying resistance to replication stress. In many cases, however, this likely reflects reduced cell replication after CRISPR-mediated inactivation of the corresponding genes under basal conditions (e.g., DLD, DLST, OGDH, Tables S2-4) and the fact that cytotoxicity induced by DHODH inhibitors or ATR inhibitors requires cell cycle progression (Fig. S1A).

To test the idea that KDM6A protects cells from replication stress-induced cell death, we inactivated KDM6A using two sgRNAs (Fig. 1H) in IDH-wildtype HOG EV cells. KDM6A knockout sensitized to BAY 2402234 (Fig. 1I), with further increases in sensitivity observed with co-knockout of the KDM6B paralog (Fig. 1J, Fig. S1G). KDM6A knockout (alone or in combination with KDM6B co-knockout) also sensitized HOG EV cells to ceralasertib (Fig. 1K–L). Dual knockout of KDM6A and KDM6B also synergized with BAY 2402234 and ceralasertib to cause DNA damage, as indicated by phospho-histone H2A.X (βH2A.X) induction (Fig. 1M). This effect was specific to drugs that induce replication stress, as KDM6A knockout did not sensitize cells to the purine synthesis inhibitor lometrexol or to the Akt inhibitor triciribine (Fig. S1H–I).

We next asked if ascorbate alleviated replication stress hypersensitivity of IDH-mutant glioma cells in a KDM6-dependent manner because ascorbate, as noted above, can counteract R2HG-dependent inhibition of 2OG-dependent dioxygenases (16), potentially including KDM6A and KDM6B enzymes. Indeed, co-knockout of KDM6A/B in IDH-mutant HOG cells abolished the cytoprotective effects of A2P during BAY 2402234 or ceralasertib treatment (Fig. 1N).

To test whether KDM6 enzymes protect against replication stress in other cell contexts, we inactivated KDM6A in NHA cells using CRISPR (Fig. 2A). KDM6A loss increased sensitivity to BAY 2402234 and ceralasertib at late passage (Fig. 2B), but not early passage (Fig. S2A), recapitulating the kinetics of replication stress sensitization by mutant IDH in these cells (15). To extend our findings to patient-derived models, we deleted KDM6A in patient-derived UTSW63 (Fig. 2C, Fig. S2B) and TS516 (Fig. 2D, Fig. S2C) IDH-wildtype glioblastoma glioma stem-like cells (GSCs). We previously showed that TS516 cells are resistant to DHODH inhibition both in vitro and in vivo (15). Loss of KDM6A sensitized both patient-derived GSC models to BAY 2402234 and ceralasertib. Furthermore, KDM6A loss sensitized TS516 orthotopic xenografts to BAY 2402234 in vivo (Fig. 2E–F). The modest effect size in this orthotopic xenograft model may reflect compensation by KDM6B. Together, these findings confirm KDM6 enzymes as key regulators of response to replication stress across engineered, patient-derived, and xenograft models of glioma.

**Figure 2.**
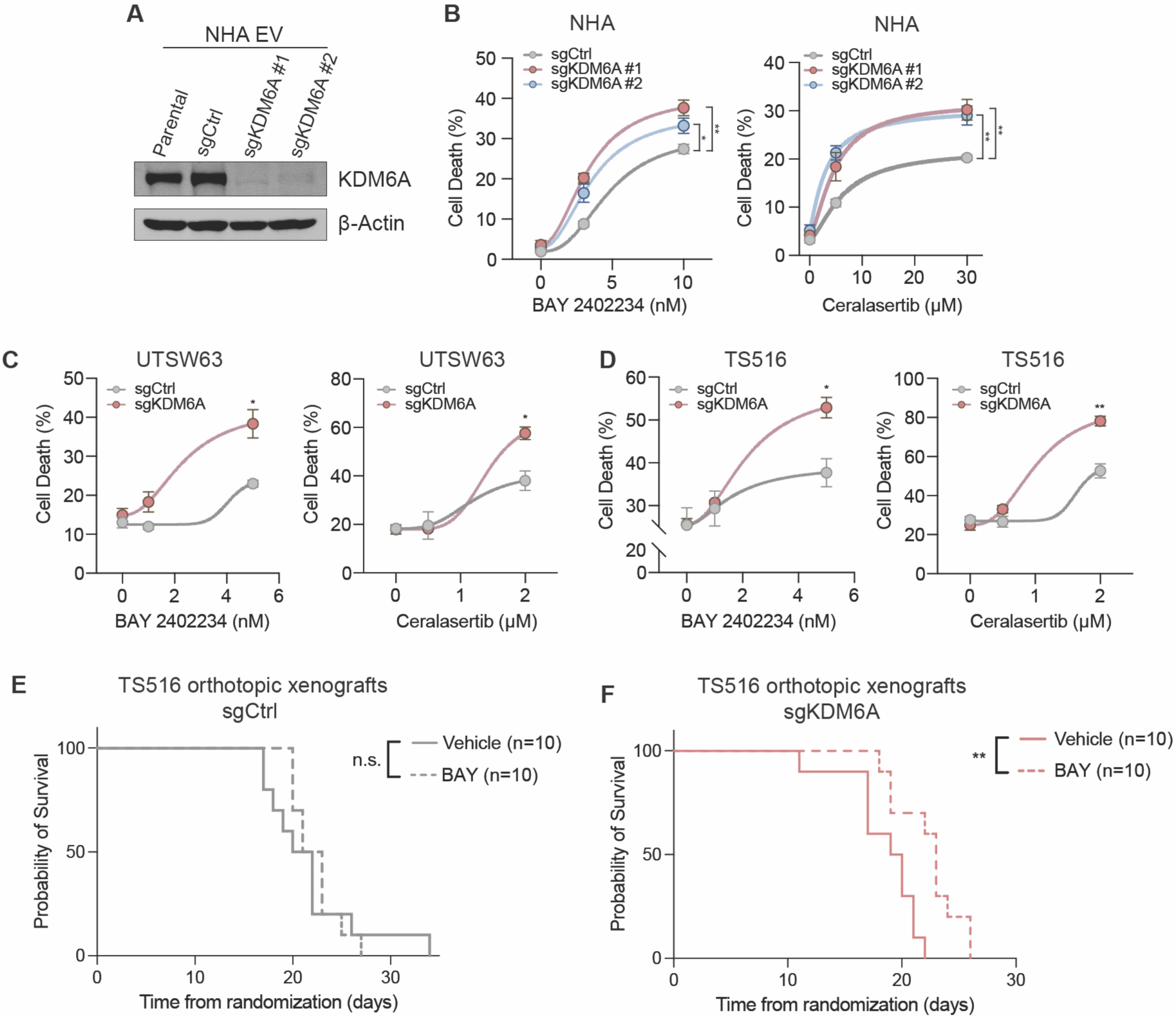
KDM6A loss sensitizes engineered and patient-derived glioma models to DHODH and ATR inhibitors. **(A)** Immunoblot analysis of NHA cells expressing an empty vector (EV), Cas9, and sgRNAs targeting KDM6A or a control sgRNA. **(B)** Cell death assay of cells in (A) (late passage) treated with BAY 2402234 or ceralasertib (*n =* 3). 10 nM BAY 2402234: sgKDM6A #1 vs. sgCtrl, p=0.0015; 10 nM BAY 2402234 sgKDM6A #2 vs. 1 sgCtrl, p=0.011; 30 µM ceralasertib: sgKDM6A #1 vs. sgCtrl, p=0.0015; 30 µM ceralasertib sgKDM6A #2 vs. 1 sgCtrl, p=0.0018. **(C–D)** Cell death assays of UTSW63 glioblastoma cells (C) or TS516 glioblastoma cells (D) with either sgKDM6A or control sgRNAs treated with BAY 2402234 or ceralasertib (*n =* 3). UTSW63: 5 nM BAY 2402234: p=0.014; 2 µM ceralasertib: p=0.015. TS516: 5 nM BAY 2402234: p=0.020; 2 µM ceralasertib: p=0.0045. **(E–F)** Kaplan-Meier survival curves of TS516 orthotopic xenografts generated from cells in (D) treated with BAY 2402234 or vehicle (*n =* 10 per cohort). TS516 sgCtrl: BAY 2402234 vs. vehicle: p=0.6892; TS516 sgKDM6A: BAY 2402234 vs. vehicle: p=0.0038. n.s. = not significant. For all panels, data are means ± SEM. Two-tailed p-values were determined by unpaired t-test. Overall survival differences were determined by log rank test.

### The demethylase activity of KDM6 mediates response to replication stress

While KDM6 enzymes canonically function as H3K27me3 demethylases, demethylase-independent functions of KDM6 have also been described and even implicated in mediating response to DNA damage (24). We thus sought to determine the specific function of KDM6 enzymes that is required for replication stress tolerance. We addressed this question by surveying interactions between functional domains of the KDM6A protein and cellular replication stress response. We performed targeted base editor mutagenesis screens using activity-based selection, which involves incorporating a defective splice-site donor site in the puromycin resistance gene that is corrected and restored upon successful base editing (25). Cell survival following puromycin treatment is thereby functionally linked to base editor activity, thus selecting cells with high editing efficiency. We used this approach with a library of sgRNAs tiling the *KDM6A* gene, non-targeting negative control sgRNAs, and pan-lethal positive control sgRNAs targeting splice donor sites of essential genes. In addition, the sgRNA library included two DHODH-targeting sgRNAs predicted to create either L358P or A58T mutations in the presence of an A>G base editor or C>T base editor, respectively (Table S5). These mutants confer resistance to BAY 2402234 (15) and were included as positive controls that would be expected to enrich in a BAY 2402234 screen.

We infected IDH-wildtype HOG EV cells with either an A>G base editor or C>T base editor, followed by this sgRNA library. We then treated cells with BAY 2402234 or DMSO and determined sgRNA abundance in each treatment group (Fig. 3A). As expected, sgRNAs causing drug-resistant DHODH L358P and DHODH A58T mutations were overwhelmingly the strongest enriched hits in the A>G and C>T base editor screens, respectively (Fig. S3A), establishing that our base editor screens were successful technically. Given that dominant “runaway hits” can mask the ability to observe other enriched or depleted sgRNAs in pooled screens, we also performed this screen using ceralasertib, where the sgRNAs encoding the L358P and A58T mutants were not expected to score. In the ceralasertib screen, we observed preferential depletion of sgRNAs encoding missense mutations in or around the JmjC catalytic domain of KDM6A compared to sgRNAs outside the JmjC domain (Fig. 3B–D, Tables S6–7). sgRNAs targeting the catalytic domain and flanking regions were depleted when comparing average z-scores (Fig. 3D) or when using a threshold cutoff (sgRNAs with z-score <-1.5, Fig. S3B). We therefore hypothesized that the JmjC domain and, by extension, the catalytic activity of KDM6A protect cells from replication stress. To empirically test this, we exogenously expressed wildtype KDM6A (KDM6A-WT), a KDM6A mutant lacking the JmjC domain (KDM6A-ΔCD), or the empty vector (EV) in IDH-wildtype HOG EV cells in which endogenous KDM6A was inactivated using CRISPR (Fig. 3E). Exogenously re-expressed KDM6A-WT, but not KDM6A-ΔCD, rescued sensitivity to BAY 2402234 and ceralasertib (Fig. 3F). We additionally exogenously re-expressed wildtype KDM6A, KDM6A-ΔCD, or EV in IDH-mutant HOG cells (Fig. S3C). Although exogenously expressed wildtype KDM6A is still expected to be inhibited by R2HG in these IDH-mutant cells, we hypothesized that increasing KDM6A expression would partially counteract inhibition by R2HG. Indeed, wildtype, but not KDM6A-ΔCD, reduced cell death induced by either BAY 2402234 or ceralasertib (Fig. S3D), indicating that the JmjC domain of KDM6A is important for replication stress response. Collectively, these results support a model in which R2HG production by mutant IDH inhibits the demethylase activity of KDM6 enzymes and thereby increases susceptibility to replication stress-dependent DNA damage (Fig. 3G).

**Figure 3.**
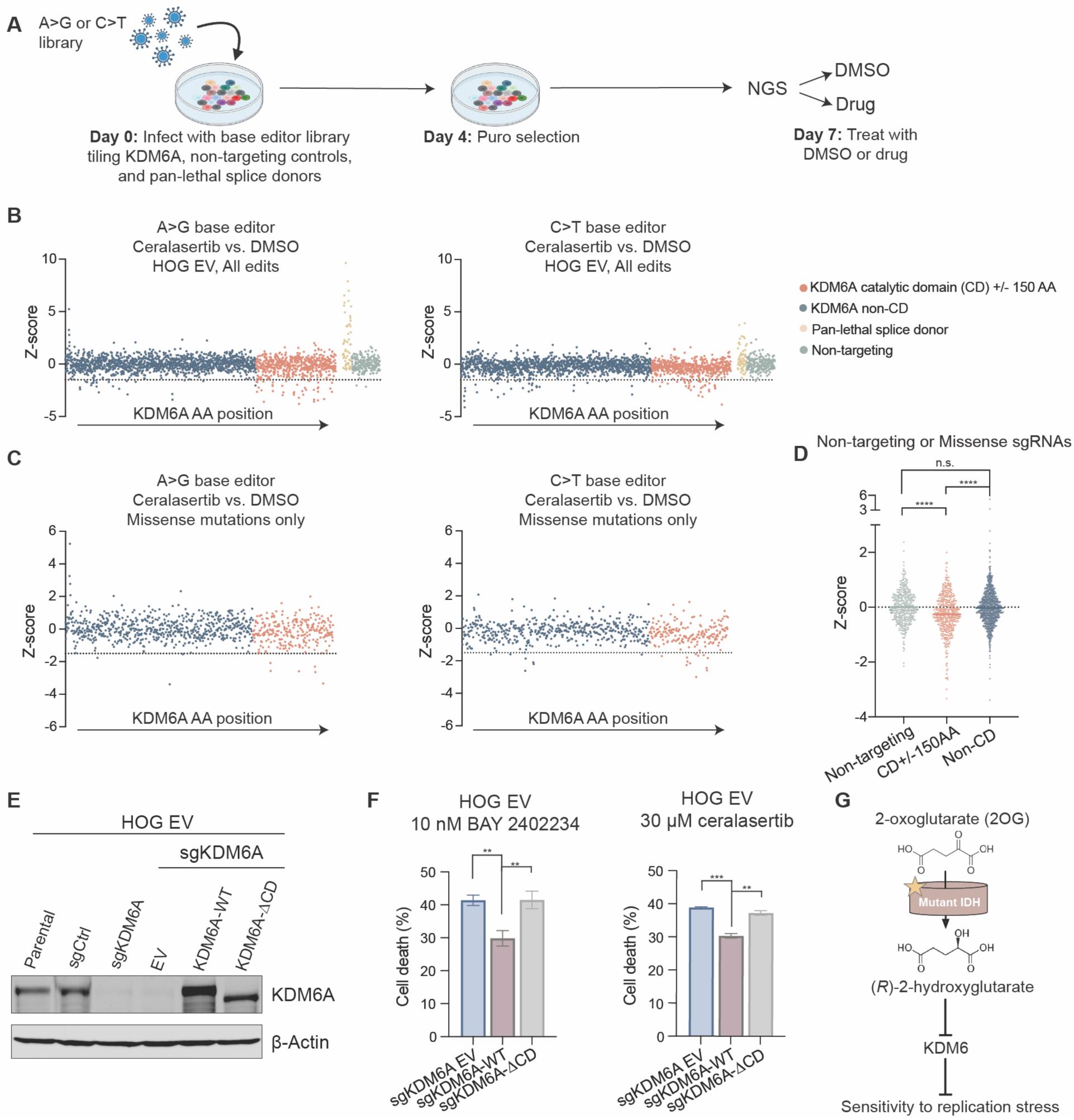
The catalytic activity of KDM6A is required for replication stress protection. **(A)** Schema of activity-based base editor screens. **(B)** Z-scores of all sgRNAs in 30 µM ceralasertib compared to DMSO treatment arms for A>G and C>T base editor screens (*n =*4 per drug per screen). **(C)** Z-scores (30 µM ceralasertib vs. DMSO) of only sgRNAs that create missense mutations in A>G and C>T base editor screens. **(D)** Average z-scores of non-targeting sgRNAs, sgRNAs creating missense mutations targeting the peri-catalytic domain (CD+/-150 AA) of KDM6A, and sgRNAs targeting regions outside of the peri-catalytic domain (Non-CD); p<0.0001. **(E)** Immunoblot analysis of KDM6A knockout HOG EV cells expressing KDM6A-WT, KDM6A-ΔCD, or EV. **(F)** Cell death assay of cells in (E) treated with BAY 2402234 or ceralasertib (*n =* 6 for sgKDM6A/WT BAY 2402234 and sgKDM6A/ΔCD BAY 2402234, *n =* 3 for all others). 10 nM BAY 2402234: EV vs. KDM6A-WT: p=0.0022; KDM6A-WT vs. KDM6A-ΔCD: p=0.0088; 30 µM ceralasertib: EV vs. KDM6A-WT: p=0.0003; KDM6A-WT vs. KDM6A-ΔCD: p=0.0021. **(G)** Schema depicting model by which mutant IDH sensitizes gliomas to replication stress. For all panels, data are means ± SEM. Two-tailed p-values were determined by unpaired t-test. n.s. = not significant.

### GLIO-1 is an on-target, blood brain barrier-penetrant DHODH inhibitor

Our prior work established a preclinical rationale to use BAY 2402234 to target mutant IDH-induced replication stress hypersensitivity in glioma. However, BAY 2402234 is no longer undergoing clinical development for cancer therapy. To identify an alternative DHODH inhibitor that could be translated to early phase clinical trials in glioma, we evaluated analogs of vidofludimus calcium. Vidofludimus calcium is a DHODH inhibitor being tested in large multicenter phase 3 trials (NCT05134441, NCT05201638) for the treatment of multiple sclerosis. Prior phase 2 testing established that this drug has an appealing safety profile (26) relative to other DHODH inhibitors (27–31). Among the vidofludimus calcium analogs tested, GLIO-1 displayed several promising properties relevant to brain tumor therapy, including robust brain penetrance, high potency, and oral bioavailability (Fig. 4A, S4A). It preferentially killed IDH-mutant HOG cells (Fig. 4B) and NHA cells (Fig. 4C) compared to EV-expressing, IDH-wildtype isogenic controls. As expected, KDM6A knockout (and KDM6A/B co-knockout), like mutant IDH, sensitized IDH-wildtype HOG cells to GLIO-1 (Fig. S4B–C). Importantly, co-treatment of IDH-mutant HOG cells with supraphysiologic levels (100 µM) of the pyrimidine salvage pathway substrate uridine completely rescued cytotoxicity induced by GLIO-1, indicating that killing by GLIO-1 is likely on-target (Fig. 4D). To formally test whether GLIO-1 acts via on-target inhibition of DHODH, we treated IDH-mutant cells expressing the drug-resistant DHODH-A58T mutant, wildtype DHODH, or an empty vector (EV), with GLIO-1. DHODH-A58T-expressing cells were fully resistant to GLIO-1 at doses ten times higher than those that preferentially kill IDH-mutant cells (Fig. 4E, Fig. 4B). Furthermore, GLIO-1 displayed similar efficacy (Fig. 4F) and superior tolerability (Fig. 4G) compared to BAY 2402234 when administered to mice bearing orthotopic, patient-derived xenografts of IDH-mutant glioma. These findings support GLIO-1 as a novel, blood brain barrier-penetrant, and on-target DHODH inhibitor that is effective in preclinical IDH-mutant glioma models.

**Figure 4.**
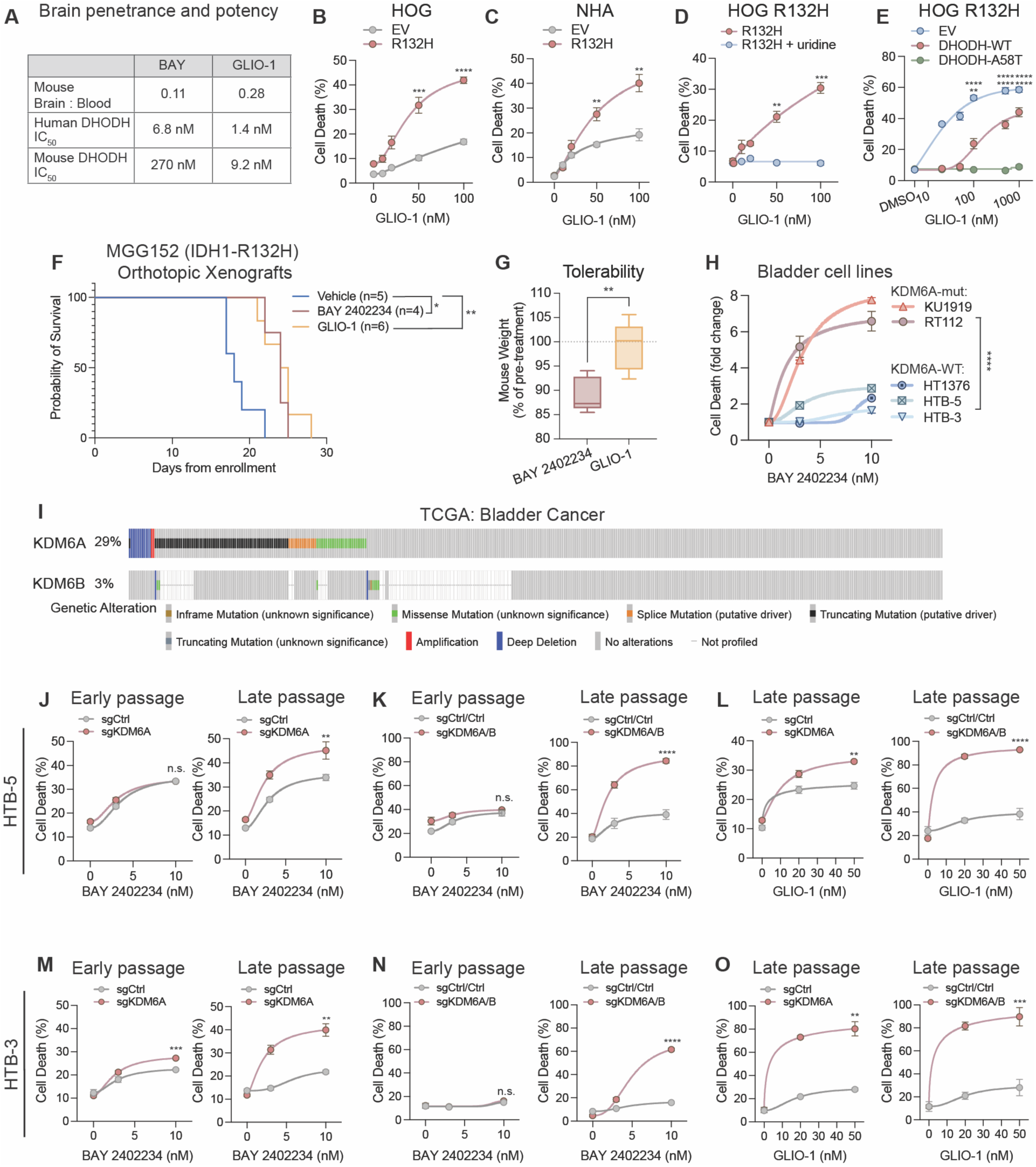
GLIO-1 is an on-target DHODH inhibitor that is effective in IDH-mutant gliomas and KDM6-mutant bladder cancers. **(A)** Brain penetrance (in mice after 2 hours P.O. dosing of 20 mg/kg) and IC_50_ against human and mouse DHODH of BAY 2402234 and GLIO-1. **(B)** Cell death assays of HOG EV or IDH1-R132H cells treated with GLIO-1 (*n =* 3). 50 nM: p=0.0004; 100 nM: p<0.0001. **(C)** Cell death assays of NHA EV or IDH1-R132H expressing cells treated with GLIO-1 (*n =* 3). 50 nM: p=0.0017; 100 nM: p=0.0011. **(D)** Cell death assays of HOG R132H cells treated with GLIO-1 in the presence or absence of 100 µM uridine supplementation (*n =* 3). 50 nM: p=0.0013, 100 nM: p=0.0002. **(E)** Cell death assays of HOG R132H cells expressing a drug-resistant point mutant of DHODH (DHODH-A58T), wildtype DHODH (DHODH-WT) or an empty vector (EV) treated with GLIO-1 (*n =* 3). 100 nM: EV vs. DHODH-A58T: p<0.0001, DHODH-WT vs. DHODH-A58T: p=0.0026; 500 nM: EV vs. DHODH-A58T: p<0.0001, DHODH-WT vs. DHODH-A58T: p<0.0001; 1000 nM: EV vs. DHODH-A58T p<0.0001, DHODH-WT vs. DHODH-A58T: p<0.0001. **(F)** Kaplan-Meier survival curves of MGG152 orthotopic xenografts treated with vehicle (*n =* 5), BAY 2402234 (*n =* 4), or GLIO-1 (*n =* 6); vehicle vs. BAY 2402234: p=0.015; vehicle vs. GLIO-1: p=0.0057. **(G)** Average weights of mice treated in (F) following 11 days of treatment with the indicated drugs, p=0.0038. **(H)** Cell death assays of endogenous KDM6A-mutant or KDM6A wildtype bladder cancer cell lines treated with BAY 2402234 for 4–5 population doublings (*n =* 3 per cell line), KDM6A-mutant vs. KDM6A-WT: p<0.0001. **(I)** Alterations in KDM6A or KDM6B in the Bladder Cancer TCGA data set (MSK/TCGA, *n =* 438). **(J)** Cell death assays of HTB-5 (KDM6A-WT) cells harboring knockout of KDM6A or control sgRNA treated with BAY 2402234 at early and late passage (*n =* 3). 10 nM (early passage): p=0.877; 10 nM (late passage): p=0.0072. **(K)** Cell death assays of HTB-5 (KDM6A-WT) cells harboring co-knockout of KDM6A/B or control sgRNAs treated with BAY 2402234 at early and late passage (*n =* 3). 10 nM (early passage): p=0.367; 10 nM (late passage): p<0.0001. **(L)** Cell death assays of late-passage HTB-5 cells harboring KDM6A knockout or KDM6A/B co-knockout and their respective control sgRNAs treated with GLIO-1 (*n =* 6 for sgCtrl/Ctrl and sgKDM6A/B DMSO and 50 nM GLIO-1, *n =* 3 for all others). 50 nM: sgKDM6A vs. sgCtrl: p=0.0028; sgKDM6A/B vs. sgCtrl/Ctrl: p<0.0001. **(M)** Cell death assays of HTB-3 (KDM6A-WT) cells harboring knockout of KDM6A or control sgRNA treated with BAY 2402234 at early and late passage (*n =* 3). 10 nM (early passage): p=0.0002; 10 nM (late passage): p=0.003. **(N)** Cell death assays of HTB-3 (KDM6A-WT) cells harboring co-knockout of KDM6A/B or control sgRNAs treated with BAY 2402234 at early and late passage (*n =* 3). 10 nM (early passage): p=0.176; 10 nM (late passage): p<0.0001. **(O)** Cell death assays of late-passage HTB-3 cells harboring KDM6A knockout or KDM6A/B co-knockout and their respective control sgRNAs treated with GLIO-1 (*n =* 3). 50 nM: sgKDM6A vs. sgCtrl: p=0.0011; sgKDM6A/B vs. sgCtrl/Ctrl: p=0.0006. For panels (B) and (C), data are means ± SD. For all other panels, data are means ± SEM. Two-tailed p-values were determined by unpaired t-test.

### Replication stress inducers, including GLIO-1, are effective in KDM6-altered bladder cancers

We next asked whether sensitization to DHODH inhibitors by KDM6 inhibition extends to non-neural cells, focusing on bladder cancer because KDM6A or KDM6B loss-of-function mutations are observed in almost one-third of patients with this disease (Fig. 4I). Bladder cancer cell lines with endogenous KDM6A mutations displayed increased sensitivity to BAY 2402234 (Fig. 4H, Fig. S4D) and ceralasertib (Fig. S4E) compared to KDM6A wildtype lines. To address the possibility that KDM6A loss may correlate with, but not cause, increased sensitivity to DHODH inhibition, we generated isogenic KDM6A or KDM6A/B knockout stable cell lines using endogenous KDM6 wildtype bladder lines (HTB-5, HTB-3, and HT1376) (Fig. S4F–G, I–J, L–M). Knockout of KDM6A, alone and in combination with KDM6B, sensitized HTB-5 cells to BAY 2402234. This sensitization occurred at late, but not early passage (Fig. 4J), consistent with the slow kinetics of epigenetic alterations observed in NHA cells. Knockout of KDM6A/B together more potently sensitized cells to BAY 2402234, similarly at late, but not early passage (Fig. 4K). The same pattern was observed in HTB-5 cells treated with GLIO-1, where KDM6A knockout alone increased sensitivity to GLIO-1 and KDM6A/B dual knockout sensitized cells further, with nearly all cells being killed by 20-50 nM GLIO-1 (Fig. 4L, Fig. S4H). Similar effects were observed with HTB-3 cells (Fig. 4M–O, Fig. S4K) and HT1376 cells (Fig. S4N–O). These results nominate KDM6A and KDM6B loss-of-function mutations as predictive biomarkers of response to DHODH inhibitors and, more generally, to replication stress-inducing therapies in bladder cancer.

## DISCUSSION

Using unbiased genetic screens and validation experiments across isogenic and patient-derived in vitro and in vivo models, we discovered that KDM6 enzymes are the relevant targets of R2HG that mechanistically link IDH mutations with replication stress hypersensitivity. We showed that the catalytic activity of KDM6A is necessary to protect cells from replication stress, strongly suggesting that R2HG-mediated inhibition of KDM6 enzymes is the mechanism by which IDH-mutant gliomas are sensitized to replication stress. These findings motivated our discovery of a new DHODH inhibitor, GLIO-1, that is effective against IDH-mutant glioma xenografts and well-tolerated in vivo. Finally, we demonstrate that KDM6-mutant bladder cancers are sensitized to DHODH inhibitors, nominating a new biomarker and drug combination that is well-positioned for clinical translation.

Several groups have linked mutant IDH with sensitivity to replication stress and clinical trials aimed at exploiting this sensitivity are in various stages of development for IDH-mutant gliomas (32). Such a strategy remains especially relevant given that mutant IDH inhibitors have limited efficacy in high-grade IDH-mutant glioma patients (33). Nonetheless, how, mechanistically, mutant IDH impairs replication stress tolerance has remained incompletely understood. *IDH* mutations induce sensitivity to PARP inhibitors, although the mechanism is debated. One group attributed this to defects in homology-direct repair (HDR) via modulation of KDM4 activity (2), while a second group linked this to increased heterochromatin independent of HDR defects (6). Regarding the former, KDM4 enzymes did not score as hits in our screens. This may be due to a specific interplay between HDR activity and PARP inhibitor response that is less relevant to drugs that act nearly exclusively in a replication stress-dependent manner. Depletion of R2HG readily reverses KDM4-mediated HDR defects but not DHODH inhibitor sensitivity (15), providing additional evidence that distinct mechanisms likely underlie response to PARP inhibitors versus DHODH or ATR inhibitors.

Our findings agree with published work demonstrating the role of H3K27 methylation in maintaining genomic stability. H3K27 methylation at replication forks facilitates replication fork restart and PARP inhibitor sensitivity via recruitment of the endonuclease MUS81 (34). In addition, inhibition of KDM6 catalytic activity decreases DNA repair gene expression and sensitizes tumors to radiation therapy, possibly due to repression of specific DNA repair genes. Others have reported that the oncohistone H3K27M, which causes loss of H3K27me3 and transcriptional derepression, also increases sensitivity to replication stress (35,36). In sum, both increased and decreased H3K27 methylation have been linked to DNA replication stress, suggesting that this critical histone mark must be finely tuned for DNA replication and DNA repair (37).

We describe a new blood brain barrier-penetrant DHODH inhibitor, GLIO-1, that is effective in models of both IDH-mutant glioma and KDM6A-mutant bladder cancer. Previous work revealed impaired DNA damage responses in KDM6A-mutated bladder cancers due to H3K27me3-driven repression of DNA repair enzymes, leading to genomic instability and improved response to immunotherapy (38). Our findings nominate KDM6 loss as a biomarker that also predicts sensitivity to DNA replication stress inducers such as DHODH inhibitors. Targeting DHODH has shown promise as a treatment strategy in other cancer types including small cell lung cancer (39), nasopharyngeal carcinoma (40), diffuse midline gliomas (41), acute myeloid leukemia (42,43), KRAS-mutant cancers (44), and myc-driven tumors (45–47), underscoring the need for effective and well-tolerated DHODH inhibitors such as GLIO-1. Taken together, our results provide new understanding of the molecular mechanism underlying replication stress sensitivity conferred by mutant IDH in glioma and provide a rationale for biomarker-driven clinical trials to test the DHODH inhibitor GLIO-1 in IDH- or KDM6-mutant cancers.

## ACKNOWLEDGEMENTS

This manuscript is subject to HHMI’s Open Access to Publications policy. HHMI lab heads have previously granted a nonexclusive CC BY 4.0 license to the public and a sublicensable license to HHMI in their research articles. Pursuant to those licenses, the author-accepted manuscript of this article can be made freely available under a CC BY4.0 license immediately upon publication.

## AUTHOR CONTRIBUTIONS

Conceptualization: DDS, SKM, WGK

Resources: JAL, JGD, DV, CG, HK, SKM, DDS, WGK, DPC, KGA

Data curation: AC-YT, MDL, VTP, DMJ, VGD, EGK, LMD, JAL, JGD, WGK, SKM, DDS, CG

Software: EGK, LMD, JGD

Formal analysis: AC-YT, MDL, DDS, LMD, EGK

Supervision: DDS, SKM, WGK, JGD

Funding acquisition: DDS, SKM, WGK, DN, HK

Validation: AC-YT, MDL, DMJ, VGD, CG, DDS, VTP

Investigation: AC-YT, MDL, DMJ, VGD, VTP, DDS, HW, JGD, KWM, CG

Visualization: DDS, SKM, AC-YT, MDL, DMJ

Methodology: SKM, DDS, JGD, WGK, CG, JAL

Writing – original draft: DDS, SKM, WGK

Project administration: DDS, SKM, WGK

Writing – review & editing: AC-YT, MDL, VTP, DMJ, VGD, EGK, LMD, JAL, JGD, WGK, KWM, DN, DV, CG, HK, KGA, HW, DPC, SKM, DDS

