## Supplementary material for "KDM6 Enzymes are the Mechanistic Targets of Mutant IDH that Dictate Replication Stress Sensitivity": Methods and Supplemental Figures

### Cell Lines and Cell Culture

All cell lines were cultured at 37ºC with ambient oxygen and 5% $CO_2$. HOG cells (human oligodendroglioma line from a male) were a gift of P. Paez, SUNY Univ. at Buffalo and were cultured in Iscove's Modified Dulbecco's Medium (IMDM, Gibco, 12440061) with 10% FBS (GeminiBio, 100-106), 1% penicillin/streptomycin (Gibco, 15140163), and 2.5 mg/mL plasmocin (InvivoGen, ant-mpt).

NHA cells (human astrocytes immortalized with HPV E6 and E7 and hTERT) were generated, selected for, and maintained as previously described (15). NHA cells were cultured in Dulbecco's Modified Eagle Medium (DMEM, Gibco, 11995073) with 10% FBS (GeminiBio, 100-106) and 1% penicillin/streptomycin (Gibco 15140163). Stable HOG and NHA lines expressing either empty vector (EV) or IDH1-R132H were generated, selected for, and maintained as previously described (15).

TS516 cells were obtained from I. Mellinghoff at MSKCC, and UTSW63 cells were obtained as previously described (15,48,49). TS516 and UTSW63 cells were cultured in Neurocult NS-A Basal Medium (Human) with Proliferation Supplement (StemCell Technologies 05750) supplemented with EGF (20 ng/mL), bFGF (20 ng/mL), heparin (2 μg/mL), 1% penicillin/streptomycin, amphotericin B (250 ng/mL) and plasmocin (2.5 μg/mL). All GSC cells were maintained as neurospheres in ultra-low-adherence culture dishes and dissociated 1–2 times per week with accutase (StemCell Technologies 07922).

HTB-3 (ATCC, SCaBER, RRID:CVCL_3599), HTB-5 (ATCC, TCCSUP, RRID:CVCL_1738), and HT1376 (RRID:CVCL_1292) cells were purchased from American

Type Culture Collection (ATCC). HTB-3, HTB-5, and HT1376 cells were cultured in Eagle's Minimum Essential Medium (ATCC, 30-2003) with 10% FBS (GeminiBio, 100-106) and 1% penicillin/streptomycin (Gibco, 15140163).

**Animals**

Animals were housed and cared for according to Good Animal Practice as defined by the US Office of Laboratory Animal Welfare and approved by the Dana-Farber Cancer Institute (protocol 04-019) or the UT Southwestern Medical Center (protocol 2019-102795) Institutional Animal Care and Use Committee. Mice were housed together (up to 5 mice per cage). Animals were provided access to standard diet and water as needed. Animal welfare assessments were carried out daily during treatment periods.

**TS516 orthotopic xenograft in vivo efficacy studies**

TS516 orthotopic xenografts were generated using high-passage isogenic sgKDM6A- or control sgRNA-expressing neurosphere cells. Neurospheres were dissociated with accutase on the day of intracranial injection and resuspended in HBSS to a concentration of 2.5E4 cells/mL. Female BL6 SCID mice aged 5-6 weeks (Jackson Laboratory 001913) were anesthetized via isoflurane (2-3%) and immobilized using a stereotactic frame. Skin above the skull was shaved and disinfected with betadine and 70% ethanol. An incision was made to expose the skull surface, and a hole was drilled into the skull at 2.0 mm right lateral and 0.5 anterior to the bregma. 3 µL of TS516 cell suspension (7.5E4 cells) in a 5 µL syringe (Hamilton) was adjusted to 3.5 mm deep to the surface of the brain and then withdrawn to 3.0 mm depth. The 3 µL cell suspension mixture was implanted over the course of 3 minutes using an autoinjector (Stoelting 53311) followed by a 1-minute incubation. The syringe was then withdrawn over the course of 1 minute. The skin was closed with 4-0 vicryl sutures, and buprenorphine was given for analgesia.

Ten days after intracranial implantation, mice were randomized to diet containing either 25 mg/kg BAY 2402234 (Research Diets D22011901Y) or vehicle diet (Research Diets D11112201). Dose of BAY 2402234 in diet was chosen to achieve a target of 4 mg/kg/day dose. Mice consumed BAY 2402234 or vehicle diets freely. Diets were replaced every week throughout study duration. Mice were monitored daily with behavioral assessment and weights. Mice were placed on treatment breaks from BAY 2402234 if body weights decreased to >10% below body weight measured at time of randomization. BAY 2402234 treatment was reinstated upon recovery of body weight. Mice were euthanized when they displayed either neurological symptoms and/or met criteria for humane endpoints as specified on IACUC protocols.

**MGG152 orthotopic xenograft in vivo efficacy and tolerability studies**

MGG152 orthotopic xenografts were generated by first dissociating cells with accutase on the day of intracranial injection, followed by resuspension in HBSS to a concentration of 3.3E4 cells/μL. Female BL6 SCID mice aged 5-6 weeks (Jackson Laboratory 001913) were anesthetized via isoflurane (2-3%) and immobilized using a stereotactic frame. Skin above the skull was shaved and disinfected with betadine. An incision was made to expose the skull surface, and a hole was drilled into the skull at 2.0 mm right lateral and 3.0 anterior to the lambda. 3 μL of MGG152 cell suspension (1E5 cells) in a 5 μL syringe (Hamilton) was adjusted to 3.0 mm deep to the surface of the brain and then withdrawn to 2.5 mm depth. The 3 μL cell suspension mixture was implanted over the course of 3 minutes followed by a 1-minute incubation. The syringe was then withdrawn over the course of 1 minute. The skin was closed with surgical clips. Extended release buprenorphine and carprofen were given for analgesia.

Mice were randomized to vehicle diet (Research Diets D11112201), diet containing 25 mg/kg BAY 2402234 (Research Diets D22011901Y), or diet containing 50 mg/kg GLIO-1 (Research Diets D25082101) 12 days following intracranial implantation. Mice were monitored daily for behavioral assessment and body weight and euthanized upon displaying neurological symptoms or any criteria meeting human endpoints per IACUC protocols.

**Chemicals**

BAY 2402234 was purchased from SelleckChem (S8847). Ceralasertib (S7693), palbociclib (S1116), cisplatin (S1166) and triciribine (S1117) were purchased from SelleckChem. Lometrexol was purchased from Sigma Aldrich (SML0040). Ascorbate 2-phosphate was purchased from Thermo Fisher (A25215G).

**Vectors**

All Gateway-based cloning was performed using the following reaction conditions: The BP reactions contained 150 ng fragment, 150 ng pDONR223, BP Clonase (Fisher, 11789100), and nuclease-free water up to 5 μL. Fragment inserts <2kb were incubated for 2 hours at RT, and fragments >2kb were incubated overnight at RT. BP reaction mixtures were transformed into chemically competent HB101 *Escherichia coli* (Promega, L2011), plated on LB-Agar plates, and incubated overnight at 37°C. Single bacterial colonies were inoculated, then had plasmid DNA extracted using the QIAprep Spin Plasmid Miniprep Kit (Qiagen, 27106). LR reactions were done with 150 ng of the destination vector, 150 ng of the pDONR clone, and LR Clonase (Fisher,

11791100), and nuclease-free water up to 5 µL. Fragment inserts <2kb were incubated for 2 hours at RT, and fragments >2kb were incubated overnight at RT. LR reaction mixtures were then transformed and extracted as per above BP reactions.

lentiCRISPR v2 sgRNA expression vector for lentiviral knockouts was obtained from Addgene (52961). Oligos corresponding to sgRNA targeting the indicated genes or controls were ordered from Integrated DNA Technologies (IDT), then cloned into the lentiCRISPR v2 vector backbone via restriction enzyme digest. To anneal the sgRNA knockout guides, the oligos were mixed in equimolar ratio in a reaction that contained 10X T4 DNA ligation buffer with ATP (New England Biolabs, M0202L). The mixture was incubated for 30 minutes at 37°C and for 5 minutes at 95°C and cooled to 25°C at a rate of 5°C/minute in a thermocycler. The annealed oligos were diluted 1:200 in nuclease-free water. The lentiCRISPR v2 backbone was first digested with FastDigest Esp3l (Thermo Fisher Scientific FD0454) for 30 minutes at 37°C. The linearized backbone was then gel-extracted and purified using a QIAquick Gel Extraction Kit (Qiagen, 28706), then cleaned using a Monarch PCR and DNA Cleanup Kit (New England Biolabs, Catalog No. T1030L). 1 µL of the annealed oligos was incubated with 50 ng of the linearized lentiCRISPR v2 backbone, 1 µL of 10X T4 DNA ligation buffer (New England Biolabs, M0202L), 1 µL of T4 DNA ligase (New England Biolabs, M0202L), and nuclease-free water (up to 11 µL). The ligation mixture was incubated at room temperature (RT) for 15 minutes. Then the ligation mixture was transformed into chemically competent HB101 *Escherichia coli* (Promega, L2011) or XL10 Gold ultracompetent cells (Agilent, 200315), plated on ampicillin-containing LB-Agar plates, and incubated overnight at 37°C. Ampicillin-resistant single bacterial colonies were inoculated and expanded in ampicillin-containing liquid lysogeny broth (LB). Plasmid DNA was extracted using the QIAprep Spin Plasmid Miniprep Kit (Qiagen, 27106) and validated via whole plasmid sequencing.

Exogenous overexpression constructs for an empty vector non-coding control sequence (EV), KDM6A-WT, and KDM6A-ΔCD were ordered as gene fragments with Gateway recombination sites from Twist Biosciences and cloned into a Ubc-pLX302 destination vector. The LR reactions contained 150 ng pLX302-Ubc destination vector, 150 ng pDONR clone, LR Clonase (Fisher, 11791100), and nuclease-free water up to 5 µL. The LR reactions were incubated at RT (2 hours for fragments ≤2 kb, overnight for fragments >2kb) before being transformed into chemically competent HB101 *Escherichia coli* (Promega, L2011), plated on LB-Agar plates, and incubated overnight at 37°C. Single bacterial colonies were picked, inoculated, then had plasmid DNA extracted using the QIAprep Spin Plasmid Miniprep Kit (Qiagen, 27106). The final Ubc-pLX302 exogenous overexpression constructs were validated via whole plasmid sequencing.

Generation of WT and DHODH A58T mutant human cDNAs and corresponding EV control lentiviral vectors (pLX304-Ubc-EV-IRES-GFP, pLX304-Ubc-DHODH-WT-IRES-GFP, and pLX304-Ubc-DHODH-A58T-IRES-GFP) was previously reported (15).

**Transfection and Transduction of Cell Lines**

NHA, HTB-3, and HT1376 isogenic KDM6A and KDM6A/B knockout cell lines were generated via lentiviral transduction. HOG isogenic KDM6A and KDM6A/B knockout cell lines used in Fig. 1I, 1J, and Fig. S4B–C were generated via lentiviral transduction. HOG isogenic KDM6A and KDM6A/B knockout cell lines used in Fig. 1N, and 3F were generated via electroporation. HTB-5 isogenic KDM6A knockout lines were generated via lentiviral transduction. HTB-5 isogenic KDM6A/B knockout lines were generated via dual-sgRNA (sgKDM6A and sgKDM6B) expressing adenoviral transduction. UTSW63 and TS516 isogenic KDM6A knockout cells were generated via adenoviral transduction using a dual-sgRNA construct encoding two sgRNAs targeting KDM6A (or two control sgRNAs). sgRNA sequences used for all knockouts are as indicated on the following table:

| Name | Sequence (5' to 3') |
|---|---|
| sgCtrl #1 | GGGGCCACTAGGGACAGGAT |
| sgCtrl #2 | TGTTAGGCAGATTCCTTATC |
| sgKDM6A #1 | TTGTTCCTGAAGGTTACTG |
| sgKDM6A #2 | TCCTTGGCTCGACAAAAGCT |
| sgKDM6B | GGTGCTAGAAGAGATCAGCC |

**Lentiviral Production and Transduction**

HEK 293T cells were first seeded in antibiotic-free DMEM containing 10% FBS, then cotransfected the following day with the desired expression vector and the packaging plasmids psPAX2 (Addgene 12260) and pMD2.G (Addgene 12259) with TransIT-X2 Dynamic Delivery System (Mirus Bio, MIR 6004). Lentiviral particle-containing media were collected at 48 and 72 hours after transfection, pooled, filtered through 0.45 µm SCFA filters (CELLTREAT, 229766), optionally concentrated with Lenti-X Concentrator (Takara Bio, 631232) as per manufacturer's instruction, aliquoted, and stored at -80°C until use.

For lentiviral transduction, all cells were seeded between 0.1-0.3E6 per well in a 6-well plate (Fisher, 08-772-1B). The next day, lentivirus particle-containing media was added, supplemented with polybrene (Santa Cruz Biotechnology, SC-134220) to a concentration of 8 µg/mL, then spun at max speed (3500 x g) at room temperature. The next day, cells were expanded, and the day after, cells were dosed with the appropriate selection marker: puromycin (InvivoGen, ant-pr-1, 2 µg/mL for HOG cells, 1 µg/mL for HTB-3 and HTB-5 cells, and 1.5 µg/mL for HT1376 cells), blasticidin (InvivoGen, ant-bl-1, 5 µg/mL for HTB-3 cells, 10 µg/mL for HT1376 cells).

**Adenoviral Production and Transduction**

pAd/PL Cas9-sgRNA adenoviral vectors (see "Vectors" above) was digested with Pac-I (New England Biolabs, R0547L) overnight at 37ºC prior to transfection. 293Ad cells were first seeded at a density of 0.5E6 in a 6-well plate in antibiotic-free DMEM containing 10% FBS, then cotransfected the following day with the Pac-I digested vector and TransIT-LT1 Transfection Reagent (Mirus Bio, MIR 2300). Two days after transfection, cells were expanded and cultured in DMEM containing 10% FBS and 1% penicillin/streptomycin. Media was changed every 2 days until 14 days after transfection. The cells were harvested and lysed via three freeze-thaw cycles. After lysis, cells were centrifuged at 2400 x g for 15 min, then the supernatant was aliquoted and frozen at -80ºC as crude adenovirus. For viral titer amplification, 293Ad cells were seeded at 3E6 in a 10 cm plate (Corning, 353003) in DMEM containing 10% FBS and 1% penicillin/streptomycin, then the following day transfected with 100 µL of the crude adenovirus. Media was changed every 2 days until approximately 80% cytopathic effect was observed. Cells were then dislodged from the plate by pipetting media across the plate surface, freeze-thaw lysed, and centrifuged. The virus particle-containing supernatant was harvested, aliquoted, then frozen at -80ºC. The amplification step was repeated 2-3 times. Amplified adenovirus was purified using the ViraBind adenovirus Miniprep Kit (Cell Biolabs, VPK-099). The final viral eluate was stored in 10% autoclaved glycerol at -80ºC until use.

Transduction of TS516 and UTSW63 GSC cells with adenovirus was performed by seeding cells in a 10 cm plate (Corning, 353003) at a density of 10E6 cells in 10 mL media. The following day, 1-50 μL of purified adenovirus encoding two sgRNAs targeting KDM6A (sgKDM6A #1 and #2) or purified adenovirus encoding control sgRNAs (sgCtrl #1 and #2) was added directly to the media. Two to seven days later, infected cells were trypsinized and FACS sorted for $mCherry^{+}$.

**KDM6A and KDM6A/B knockouts via nucleofection**

KDM6A/B co-knockout of HOG cells in Fig.1N and KDM6A single knockouts (Fig. 3F) were performed as previously described (50). Briefly, each Cas9-RNP reaction was set up containing 1.2 μL Alt-R™ CRISPR-Cas9 sgRNA (IDT), 1.4 μL Alt-R™ CRISPR-Cas9 Electroporation Enhancer (IDT), 1.7 μL Alt-R™ S.p. Cas9 Nuclease (IDT, 1081058), and 0.7 μL sterile PBS (Life Technologies, 14190250). Cas9-RNP reactions were incubated at RT for at least 10 minutes. During incubation, cells were split, counted, and washed in sterile PBS. After incubation, 1E5 of PBS-washed cells were resuspended in 20 μL of SF-nucleofection buffer (Lonza, V4XC-2032). The cell resuspension was then mixed with the Cas9-RNP reaction and transferred to an individual well on a Nucleocuvette strip (Lonza, V4XC-2032). The mixture was electroporated by using a 4D-Nucleofector X unit (Lonza, AAF-1003X) with the electroporation condition (DN-100 for HOG cells). Immediately after electroporation, 80 μL of pre-warmed media was added to each electroporation mixture. 10 minutes after electroporation, electroporation mixtures were then transferred to individual wells of a 6-well plate (Fisher Scientific, 08-772-1B), each well containing 2 mL of pre-warmed media. The following morning, media was changed, and two days after changing media, cells were expanded to 10 cm dishes (Corning, 353003).

**Palbociclib rescue experiments**

HOG-EV or HOG-R132H cells were treated for 24 hrs with 1 μM palbociclib or DMSO in 10 cm dishes. Next, cells were split and seeded at a density of 1E5 cells per well in 6-well plates (Fisher Scientific, 08-772-1B) in media supplemented with 1 μM palbociclib or DMSO. 24 hours later, cells were dosed with either 10 nM BAY 2402234 or ceralasertib. Cells were harvested 72 hours (for BAY 2402234) or 24 hours (for ceralasertib) later for cell death quantification.

**Uridine rescue experiments**

HOG-R132H cells were seeded at a density of 1E5 cells per well in 6-well plates (Fisher Scientific, 08-772-1B) with or without 100 μM uridine supplementation. 24 hours later, GLIO-1 was added at the indicated concentrations, and cell death was quantified after 72 hours of treatment.

**Cell growth quantifications**

Cells were trypsinized with 0.25% trypsin-EDTA (Gibco, 25200114), collected, then quenched in fresh cell culture media for the appropriate cell line. After centrifugation, the trypsin-media supernatant was aspirated, and the cells were resuspended in fresh cell culture media. The cell suspension's concentration was then determined using a Vi-CELL™ XR Cell Viability Analyzer (Beckman Coulter), from which the total cell count was determined.

**In vitro DHODH inhibitor and ATR inhibitor viability assays**

Cells were seeded at a density between 1-3E5 per well in a 6-well plate (Fisher Scientific, 08-772-1B). The following day, cells were dosed with the DHODH inhibitor BAY 2402234 or the ATR inhibitor Ceralasertib (Selleckchem, S7693-25mg). Glioma cells (HOG, NHA, TS516, UTS63) and bladder cells (HTB-3, HTB-5, HT1375, KU1919, and RT112) were treated for 4-5 doublings for BAY 2402234 (Selleckchem, S8847) or GLIO-1 and 1–2 doublings for ceralasertib (Selleckchem, S7693-25mg). Early passage cell death assays were performed 1–2 passages after knockout of the indicated genes. Late passage cell death assays were performed 12–16 passages after knockout of the indicated genes.

To quantify cell death following drug treatment, cells were trypsinized with 0.25% trypsin-EDTA (Gibco, 25200114), quenched in fresh cell culture media, then collected in tubes (Corning, 352052). Cells were centrifuged and washed in PBS (Life Technologies, 14190250). Cells were again centrifuged, and excess PBS was aspirated. Cells were stained with Annexin V (Annexin V-FITC, BD Biosciences, 556420; Annexin V-PE, BD Biosciences, 556422; Annexin V-APC, BioLegend, 640920) as per manufacturer's instructions, and DAPI (Cell Signaling Technology, 4083S) at a final concentration of 1 µg/mL to identify early and late apoptotic cells, respectively. Cells were analyzed on an LSRFortessa™ (BD Biosciences) flow cytometer, and data were processed using FlowJo software (BD Biosciences). Cells that were AnnexinV$^{+}$/DAPI$^{-}$, AnnexinV $^{-}$/DAPI$^{+}$, or AnnexinV$^{+}$/DAPI$^{+}$ were considered dead, while AnnexinV $^{-}$/DAPI$^{-}$ cells were considered live.

**In vitro DHODH inhibitor and ATR inhibitor ascorbate 2-phosphate rescue experiments**

Ascorbate 2-phosphate (A2P) rescue experiments in HOG cells were performed following at least 24 hours of culture with 200 μM A2P supplemented in media. Short-term A2P experiments in NHA cells (Fig. S1E) were performed after 24 hours of 200 μM A2P treatment. Long-term A2P experiments in NHA cells were conducted after 6–8 weeks of culture with 200 μM A2P-supplemented media prior to plating for drug treatment assays.

**EdU+ assays**

Cells were plated in 6-well plates at least 24 hours before assessing Edu+ uptake. EdU (Sigma, 900584) was added to seeded cells at a final concentration of 20 μM. Following a 20-minute incubation at 37 °C, cells were washed with cold PBS, trypsinized, collected, and pelleted by centrifugation. Following an additional PBS wash, cells were then fixed by adding 100 μL 4% formaldehyde to each sample, washed, and then permeabilized with 0.2% Triton X-100 (Sigma, 648463). After an additional PBS wash, cells were incubated with 150 μL of freshly made click reaction solution (100 mM pH 7.4 Tris-HCl, 5 μM sulfo-cyanine3 azide dye [Lumiprobe, A1330], 10 mM ascorbic acid [Sigma, A92902], 2 mM copper(II) sulfate pentahydrate [Sigma, 209198]) for 30 minutes to label incorporated EdU. Finally, cells were washed with PBS, stained with Hoechst dye (Life Technologies, H3570), and analyzed by flow cytometry using appropriate PE and DAPI channels.

**ROS quantification in vitro assay**

HOG-R132H cells with or without 200 μM A2P were seeded in 6-well plates at a density of 1E5 cells per well. The following drugs were then added to 200 μM A2P or control wells: DMSO, 10 nM BAY 2402234, 30 μM ceralasertib, or 50 μM cisplatin. Cells were treated with BAY

2402234 and cisplatin for 24 hours, or 2 hours for ceralasertib and harvested for ROS quantification. Cells were trypsinized and centrifuged at 3000 g for 3 minutes at RT. The supernatant was aspirated and the pelleted cells were washed with PBS. The tubes were then centrifuged again at 3000 g for 3 minutes to pellet cells.

PBS was aspirated, and 1 µL of 5 mM CM-H2DCFA stain (Invitrogen C6827) was added. Cells were resuspended in this solution. Cells were then incubated away from light for 1 hour at room temperature. Following incubation, the tubes were centrifuged at 3000 g for 3 minutes to pellet cells. The PBS and CM-H2DCFA were aspirated off the pelleted cells. The cells were then resuspended in 3 mL of warm IMDM media, then incubated away from light for 30 minutes at room temperature.

Following incubation, the tubes were centrifuged at 3000 g for 3 minutes, then washed with 3 mL PBS. Following aspiration of the PBS wash, 100 µL of DAPI stain (Cell Signaling Technology 4083S at 5 mg/mL) was added to each sample. The cells were then resuspended and analyzed via flow cytometry to determine viable ($DAPI^-$) $FITC^+$ ($CM\text{-}H2DCFA^+$) cells.

**Doubling Time Calculations**

Doubling times of all cell lines were calculated by plating a known quantity of cells on day 0 ($N_i$), then quantifying cells 3-4 days later ($N_f$) using a Vi-CELL™ XR Cell Viability Analyzer (Beckman Coulter). Doubling time ($T_d$) was calculated using the following formula:

$$T_d = \frac{days \times \log(2)}{\log(N_f) - \log(N_i)}$$

**2-oxoglutarate dependent enzyme CRISPR screens**

For 2OG-dependent enzyme CRISPR knockout screens, 18E6 HOG EV cells expressing pLenti-Cas9-GFP (Addgene 86145) were seeded into 6-well plates (0.5E6 per well). 24 hours later, media was replaced with media containing polybrene (at a concentration of 8 μg/mL) and a sgRNA lentiviral library targeting 2OG-dependent enzymes (obtained from the Broad Institute Genetic Perturbation Platform, CP1347). Cell number and volume of lentivirus was chosen to achieve a representation of ≥5,000 cells per sgRNA and an MOI of 0.3. Following addition of polybrene and lentiviral library, cells were centrifuged at 3,500 g at room temperature for 30 minutes. After overnight incubation, cells were trypsinized and expanded and seeded at a density of 5E6 per 15 cm tissue culture dish. The following day, puromycin was added to a final concentration of 2 μg/mL. Cells were maintained in culture and passaged every 3-4 days. At the indicated time points (7 days, 16 days, and 28 days) following sgRNA library infection, cells were trypsinized and plated at a density of 1.5E6 per 15 cm tissue culture dish for drug or DMSO treatments. Remaining cells at each of these time points were pelleted for assessment of sgRNA enrichment/depletion prior to drug or DMSO treatment (Fig. S1F). For BAY 2402234 screens, 10 nM BAY 2402234 or DMSO was dosed one day after seeding. Cells were scrape harvested and flash frozen in liquid nitrogen 72 hours after dosing. For ceralasertib screens, 30 μM ceralasertib or DMSO was dosed 3 days after seeding, scrape harvested and flash frozen in liquid nitrogen 24 hours after dosing. Genomic DNA was extracted from the frozen cell pellets using a Qiagen QIAamp Blood Kit (Qiagen, 51194), then cleaned using the OneStep PCR Inhibitor Removal Kit (Zymo Research, D6030). Purified genomic DNA was then quantified using Qubit 1X dsDNA HS Assay Kit (Invitrogen, Q33230) on a Qubit 4 Fluorometer (Invitrogen, Q33238). The purified genomic DNA was then submitted to the Genetic Perturbation Platform (Broad Institute) for next-generation sequencing (NGS).

**Activity-based base Editor Screens**

HOG EV cells were infected with lentivirus generated from pRDA_867 (Addgene 231109**)** or pRDB_092 (Addgene 231110) to express either the A>G Base Editor or the C>T Base Editor, respectively. HOG EV pRDA_867 and HOG EV pRDB_092 cells (3.6E7 each) were then transduced (MOI: 40-50%) with their respective custom base editor libraries (Broad Institute, CP2130 and CP2131) targeting the full-length of *KDM6A* coding regions (representation: 2,500 cells/sgRNA). Cells were seeded in 6-well plates (Fisher Scientific, 08-772-1B) at a density of 5E5 in 1 mL of IMDM cell culture media (containing 10% FBS, 1% penicillin/streptomycin, and 2.5 mg/mL plasmocin). The following day, an additional 1 mL of cell culture media supplemented with polybrene and the respective base editor viruses was added (achieving a final volume of 2 mL and final concentration of 8 μg/mL polybrene per well). Cells were then centrifuged at 3,500 g at room temperature for 30 minutes. The following day, media was aspirated and replaced with 2 mL of fresh cell culture media. The next day, cells were trypsinized and expanded. Two days after expansion, cells were dosed with puromycin (InvivoGen, ant-pr-1) at a concentration of 2 μg/mL. Two days later, after selection was complete, cells were trypsinized, expanded, and seeded at a density of 1.5E6 per 15 cm plate for each arm, maintaining representation of 2,500 cells/sgRNA total. For BAY 2402234 screens, plates were dosed with either DMSO or 10 nM BAY 2402234. Three days after BAY 2402234 or DMSO dosing, cells were scrape harvested and flash frozen in liquid nitrogen. For ceralasertib screens, plates were dosed with either DMSO or 30 μM ceralasertib 3 days after seeding. One day after ceralasertib or DMSO dosing, cells were scrape harvested and flash frozen in liquid nitrogen.

Genomic DNA was extracted from the frozen cell pellets using a Qiagen QIAamp Blood Kit (Qiagen, 51194), then cleaned using the OneStep PCR Inhibitor Removal Kit (Zymo Research, D6030). Purified genomic DNA was then quantified using Qubit 1X dsDNA HS Assay Kit (Invitrogen, Q33230) on a Qubit 4 Fluorometer (Invitrogen, Q33238). The purified genomic DNA was then submitted to the Genetic Perturbation Platform (Broad Institute) for NGS. Analysis of activity-based base editor screen results was performed as previously described (25), including self-editing correction and log-normalization of the resultant data set.

**Validation of CRISPR-mediated KDM6B Knockout**

Genomic DNA was extracted using the QIAamp DNA Blood Mini Kit (Qiagen, Catalog No. 51104) from cell pellets of all KDM6B knockout cells and their respective controls. PCR was performed to amplify amplicons centered on the KDM6B sgRNA target site. PCRs were generated with the KOD Xtreme Hot Start DNA Polymerase kit (Fisher Scientific, Catalog No. 719753) and PCR conditions of 94ºC for 2 minutes; 35 cycles of 98ºC for 10 seconds, lowest primer $T_m$ for 30 seconds, and 68ºC for 1 minute/kb plus 3 seconds of buffer; and 10ºC for 5 minutes. Amplicons were prepared via nested PCR reactions to amplify ~280 bp regions surrounding the KDM6B sgRNA target site. The PCR products were run on a 1% agarose gel, the appropriate band was excised, and DNA was extracted using the QiaQuick Gel Extraction Kit (Qiagen, Catalog No. 28706). The gel-extracted PCR products were cleaned using the Monarch PCR and DNA Cleanup Kit (New England Biolabs, Catalog No. T1030L). Amplicons were sequenced via NGS by the Massachusetts General Hospital Center for Computational and Integrative Biology DNA Core. Crispresso2 (http://crispresso2.pinellolab.org/;

RRID:SCR_024503) was used to analyze the resulting fastq files to quantify the degree of KDM6B editing.

| Primer Name | Primer Sequence | Annealing Temperature Used |
|---|---|---|
| sgKDM6B Forward Primer | ATGACCCCCACCCAACCGCC | 61.4ºC |
| sgKDM6B Reverse Primer | GCCGCCGATGCTCCTTCTGA | |

**Immunoblot Assays of Protein Expression**

Cells were pelleted and subsequently lysed in RIPA lysis buffer (Life Technologies, 89900) containing a protease inhibitor cocktail (Sigma, 11836170001) and a phosphatase inhibitor cocktail (Sigma, 4906837001). Cell lysates were incubated on a rotor for 30 minutes at 4ºC, then centrifuged at 17,000 x g for 10 minutes at 4ºC. The supernatant was then extracted and transferred to Eppendorf tubes. Clarified lysates were quantified using the Bradford protein assay (Bio-Rad Laboratories, 5000006). Samples were prepared by mixing protein with Laemmli SDS-Sample Buffer (Boston Bioproducts, BP-111R-25ML) and RIPA lysis buffer (containing protease inhibitor and phosphatase inhibitor) to yield a final concentration of 2 µg/mL protein. Samples were boiled at 95ºC for 5 min, before resolving on Novex™ Tris-Glycine protein gels. For semi-dry transfers, lysates were transferred to nitrocellulose membranes using the Trans-Blot Turbo RTA Midi 0.2 µm Nitrocellulose Transfer Kit (Bio-Rad Laboratories, 1704271) with the Bio-Rad Trans-Blot Turbo Transfer System (Bio-Rad, 1704155). For wet-transfer, lysates were transferred to PVDF membranes (Life Technologies, 88518) overnight. Membranes were briefly stained with Ponceau S (Cell Signaling Technology, 59803) to ensure equal loading and transfer. Membranes were washed with water, blocked in Intercept Blocking Buffer (LI-COR, NC1660562) for 30 min, then incubated in primary antibody solutions prepared in Intercept

Blocking Buffer containing 0.2% Tween 20 (Sigma, P1379-25ML) overnight. The following day, blots were washed in TBST, then incubated in their respective secondary antibodies (LICOR NC9744100, LICOR NC9523609) for 1 hour. The membranes were imaged using the Odyssey Classic Imager (LI-COR, 9120), then analyzed using the Image Studio software (LI-COR, Version 5.2.5). All antibodies and their dilution factors are reported in the below table.

| Antibody Name | Manufacturer | Catalog Number | Dilution Factor |
|---|---|---|---|
| KDM6A (UTX) | CST | 33510S | 1:500 |
| γh2ax | CST | 9718S | 1:1000 |
| α-tubulin | Sigma Aldrich | T5168-100UL | 1:4000 |
| β-actin (mouse) | CST | 3700S | 1:1000 |
| β-actin (rabbit) | CST | 4970S | 1:1000 |
| Vinculin | Sigma Aldrich | V9131-100UL | 1:10000 |
| IRDye 800CW Donkey anti-Mouse IgG Secondary Antibody | LI-COR | NC9744100 | 1:10000 |
| IRDye 800CW Donkey anti-Rabbit IgG Secondary Antibody | LI-COR | NC9523609 | 1:10000 |

**Human and mouse DHODH inhibition assay**

Inhibition of human and mouse DHODH was measured in vitro using an *N*-terminally truncated recombinant human or mouse DHODH enzyme, respectively. The final assay mixture contained 60 µM 2,6-dichloroindophenol, 50 µM decylubiquinone, 100 µM dihydroorotate and the DHODH protein whose concentration was adjusted in a way that an average slope of approx. 0.2

AU/minute served as the positive control (no inhibitor). Measurements were performed in 50 mM TrisHCl, 150 mM KCl and 0.1% Triton X-100 at pH 8.0 and at 30°C with at least six different concentrations of a test compound. The reaction was started by adding dihydroorotate and measuring the absorption at 600 nm for 2 minutes. Each test compound concentration used for $IC_{50}$ calculation was tested in at least three independent experiments.

**Mouse blood and brain exposure**

The brain exposure of GLIO-1 and BAY 2402234 was evaluated in 3 female mice (C57BL/6J, 8 week old) after oral cassette dosing. All experimental procedures were approved by and conducted in accordance with the regulations of the local Animal Welfare authorities (Landesamt für Gesundheit und Verbraucherschutz, Abteilung Lebensmittel- und Veterinärwesen, Saarbrücken, Germany). Dose was 20 mg/kg, application volume was 5 mL/kg and vehicle was 5% Solutol, 95% NaCl solution (at 0.9% saline concentration). At 2 hours after dosing, 20 µL whole blood from the retrobulbar venous plexus under short isoflurane anesthesia was collected into Li-heparin tubes, frozen on dry ice within 1-2 minutes of sampling and stored at –20°C until processed for LC-MS analysis. Animals were sacrificed and perfused with PBS until PBS was transparent and brain was collected and stored at –20°C until processed for LC-MS analysis and ELISA assay for determination of albumin (the measured amounts of albumin in the brain fraction were negligible).

**Quantification and statistical analysis**

Information related to data presentation and statistical analysis for individual experiments can be found in the corresponding figure legends. Statistical analyses were carried out using GraphPad

Prism software. Significance of comparisons involving two groups was calculated by unpaired two-tailed *t*-test. For comparisons of two groups with significantly different variances, Welch's *t*-test was used. For comparisons of two groups without significant differences in variances, Student's *t*-test was used. Significance of comparisons involving three or more groups was calculated by one-way ANOVA. Significance of survival data was determined by log-rank test. Statistical evaluation of patterns of co-occurrence or mutual exclusivity was performed using one-sided Fisher's exact tests. Chi-square tests were used to determine differences between categorical groups. For all tests, p-values <0.05 were considered statistically significant.

**SUPPLEMENTAL INFORMATION**

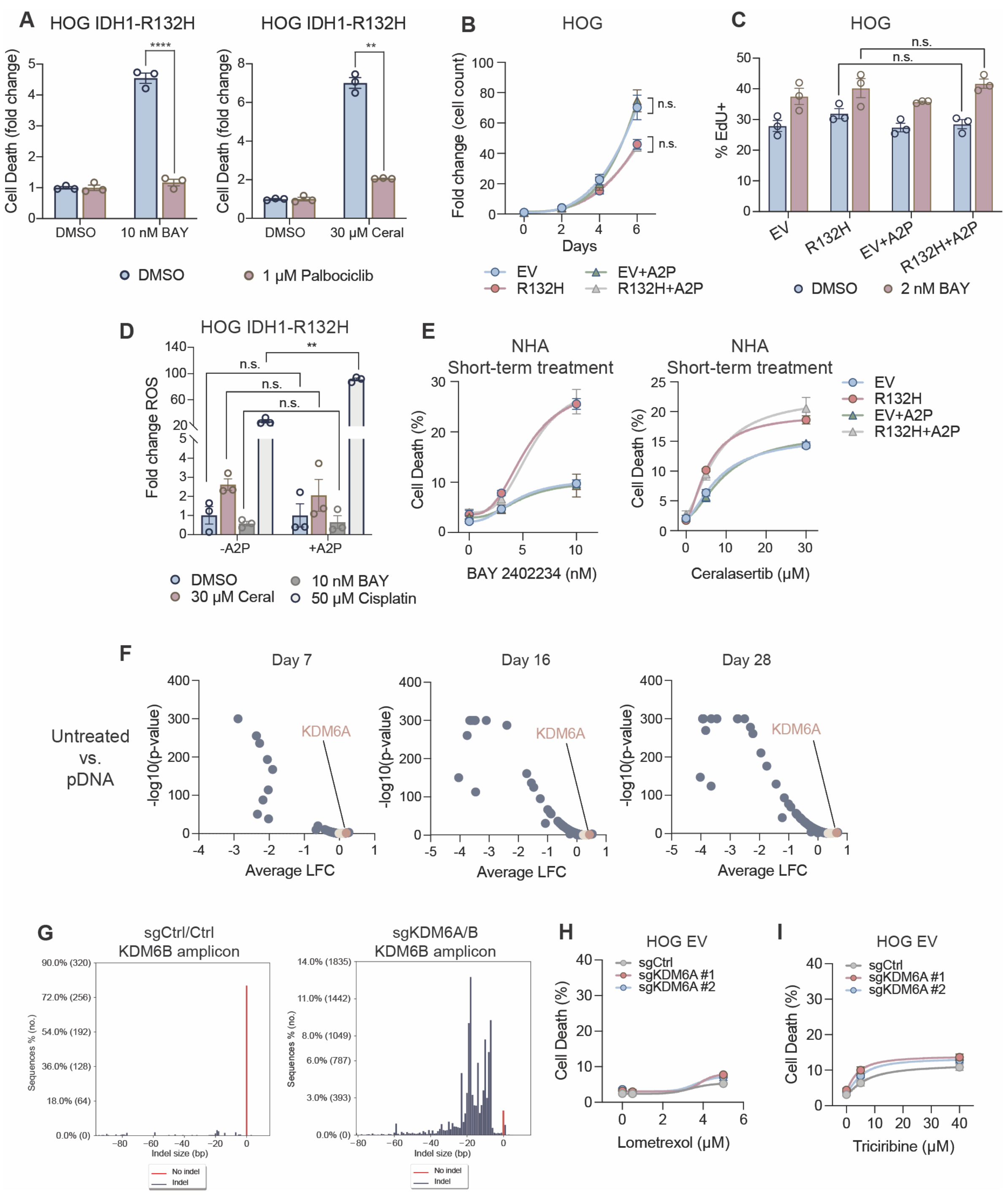


**Figure S1. Ascorbate 2-phosphate profiling studies and CRISPR screen validation experiments, related to Figure 1. (A)** Cell death assays of IDH-mutant HOG cells treated with

10 nM BAY 2402234 or 30 μM ceralasertib with or without 1 μM palbociclib ($n$ = 3). BAY 2402234/DMSO vs. BAY 2402234/palbociclib: p<0.0001; ceralasertib/DMSO vs. ceralasertib/palbociclib: p=0.0031. **(B)** Cell counts of HOG EV or IDH1-R132H (R132H) cells at the indicated days post-seeding with or without 200 μM ascorbate 2-phosphate (A2P) ($n$ = 3). **(C)** Percent EdU+ HOG EV or R132H cells treated with or without 200 μM A2P and either DMSO or 2 nM BAY 2402234. **(D)** Changes in reactive oxygen species (ROS) in HOG R132H cells treated with the indicated drugs with or without 200 μM A2P ($n$ = 3); 50 μM cisplatin vs. 50 μM cisplatin + A2P: p<0.0001. **(E)** Cell death assays of NHA EV or R132H cells treated with either BAY 2402234 or ceralasertib, with or without short-term treatment of 200 μM A2P. **(F)** Volcano plots of log fold changes (LFC) of sgRNAs in HOG EV cells harvested at the indicated days following library infection vs. library plasmid DNA (pDNA). **(G)** Indel size distribution of amplicons encompassing the target region of the KDM6B sgRNA in HOG EV cells expressing KDM6A/B sgRNAs or control sgRNAs. **(H)** Cell death assays of HOG EV cells with sgRNAs targeting KDM6A or control treated with lometrexol or triciribine ($n$ = 3). n.s. = not significant. For all panels, data are means ± SEM. Two-tailed p-values were determined by unpaired t-test.

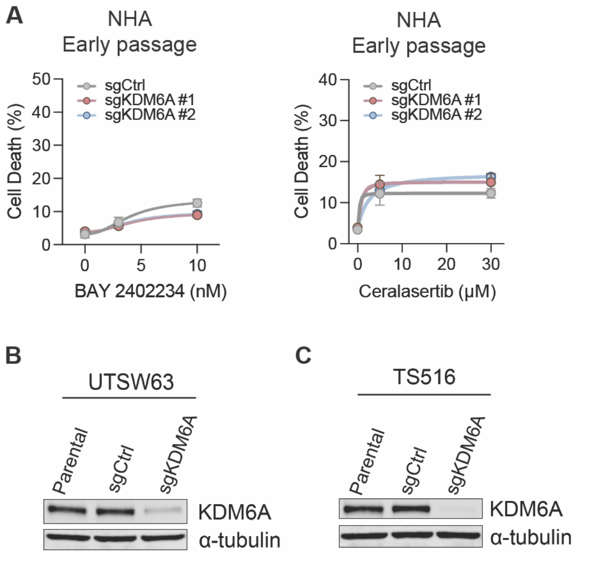


**Figure S2. Early-passage cell death assays in sgKDM6A NHA cells and validation of KDM6A knockout, related to Fig. 2. (A)** Cell death assays of NHA cells in Fig. 2A treated with BAY 2402234 or ceralasertib at early passage following knockout of KDM6A (*n* = 3). **(B–C)** Immunoblot analysis of UTSW63 cells in Fig. 2C (B) and TS516 cells in Fig. 2D–F (C). For all panels, data are means ± SEM. Two-tailed p-values were determined by unpaired t-test.

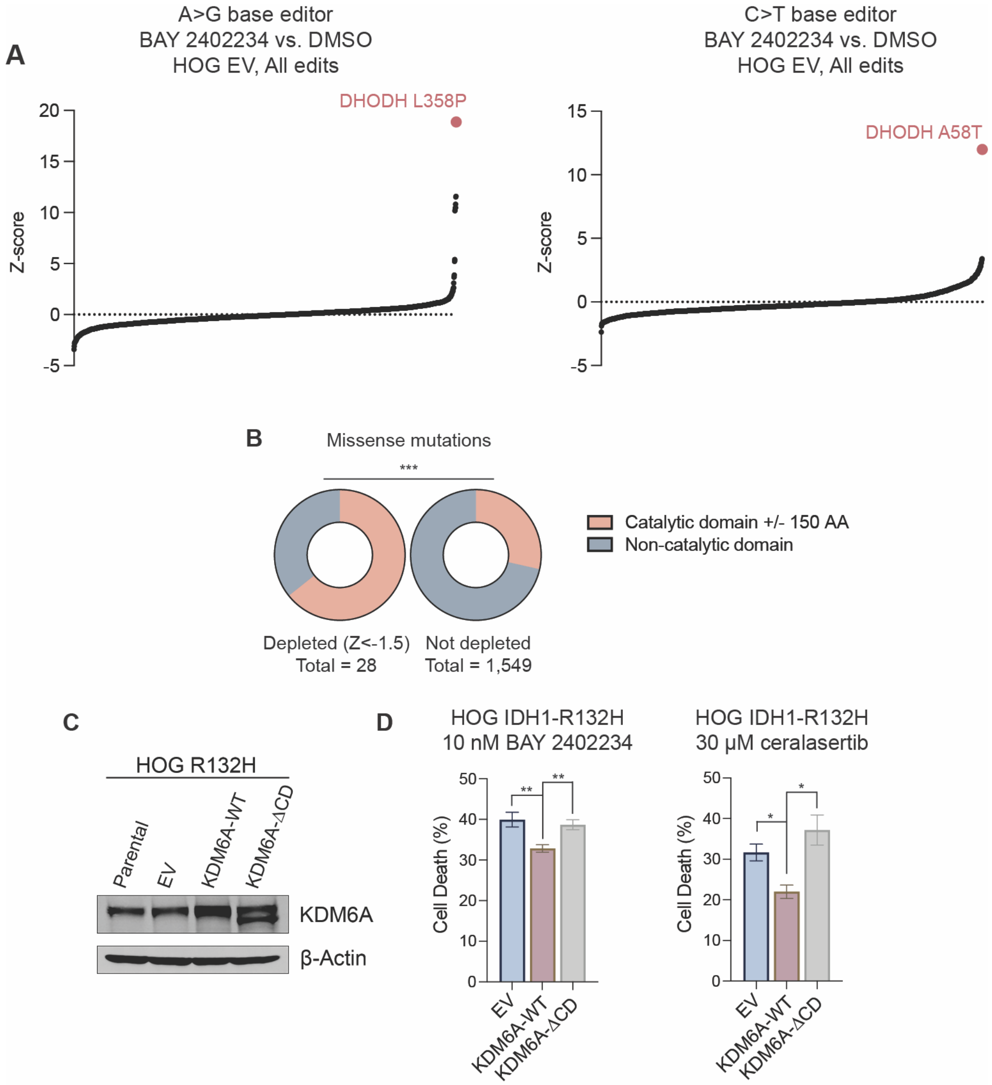


**Figure S3. Base editor screens and exogenous KDM6A rescue experiments, related to Fig. 3.** **(A)** Waterfall plot of sgRNAs in A>G and C>T base editor screens in HOG EV cells treated with 10 nM BAY 2402234 compared to DMSO control (A>G screen, $n$ = 2; C>T screen, $n$ = 3). **(B)** Distribution of sgRNAs with z-scores <-1.5 that target the peri-catalytic domain or regions outside of the peri-catalytic domain in base editor screens in Fig. 3B–D; p=0.0002. **(C)** Immunoblot analysis of HOG R132H cells expressing KDM6A wildtype (KDM6A-WT), KDM6A with a Jumonji C domain deletion (KDM6A-ΔCD), or an empty vector (EV). **(D)** Cell death assay of cells in (C) treated with the indicated doses of BAY 2402234 or ceralasertib ($n$ = 6 for BAY 2402234 and $n$ = 3 for ceralasertib). 10 nM BAY 2402234: EV vs. KDM6A-WT:

p=0.0091; KDM6A-WT vs. KDM6A-ΔCD: p=0.0041; 30 μM ceralasertib: EV vs. KDM6A-WT: p=0.024; KDM6A-WT vs. KDM6A-ΔCD: p=0.038. For all panels, data are means ± SEM. Two-tailed p-values in (D) were determined by unpaired t-test. Two-tailed p-values in (B) were determined by Fisher's exact test.

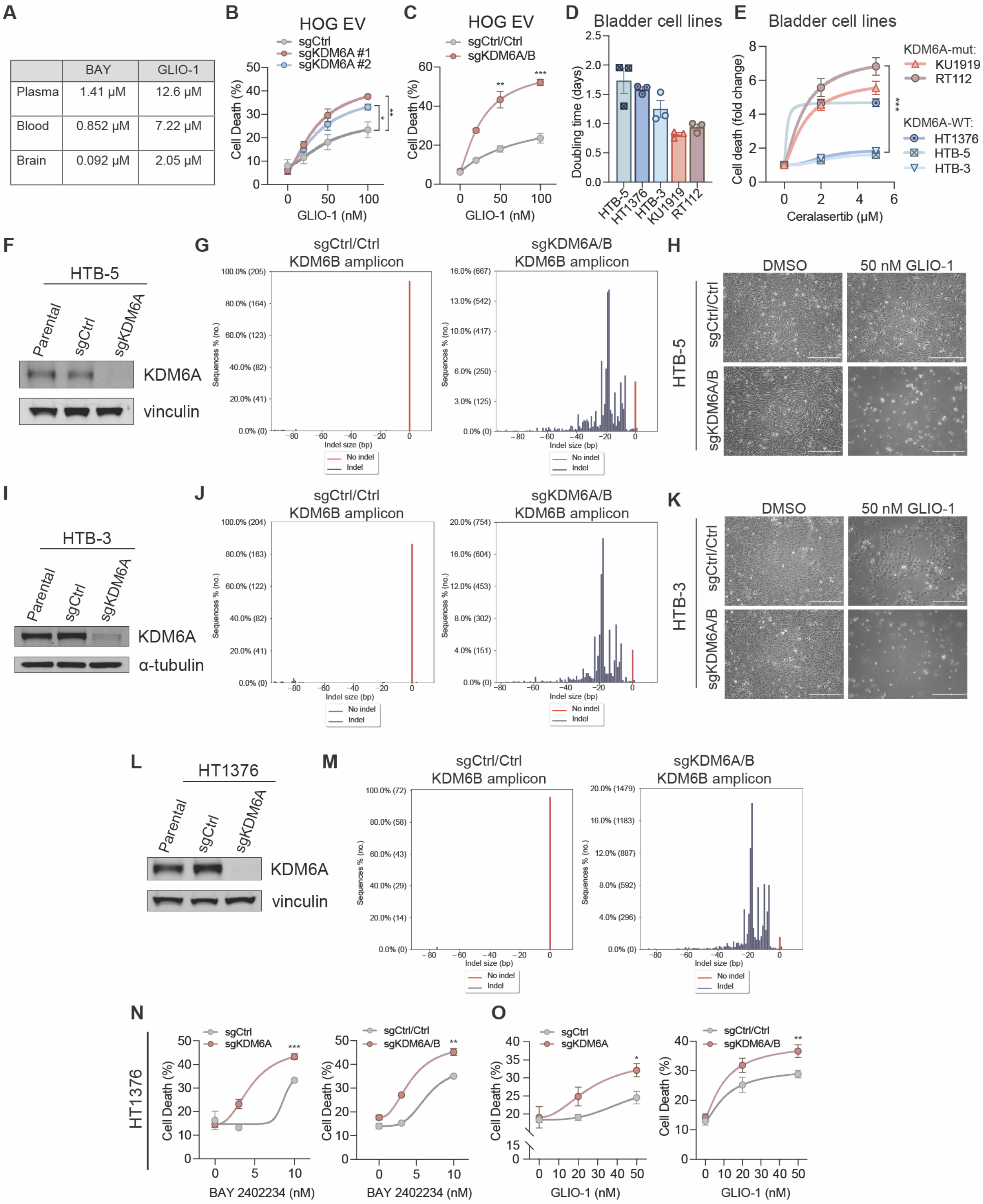


**Figure S4. Evaluation of GLIO-1 in KDM6 knockout lines and validation of KDM6A/B knockout, related to Fig. 4. (A)** Plasma, blood, and brain concentrations of BAY 2402234 and

GLIO-1 determined 2 hours after oral dosing. Values were used to calculate brain : blood ratios in Figure 4A. **(B)** Cell death assays of sgKDM6A or sgCtrl HOG EV cells treated with GLIO-1 (*n* = 3). 50 nM: sgKDM6A #1 vs. sgCtrl, p=0.0093; 50 nM sgKDM6A #2 vs. 1 sgCtrl, p=0.0318; 100 nM: sgKDM6A #1 vs. sgCtrl, p=0.0021; 100 nM sgKDM6A #2 vs. 1 sgCtrl, p=0.0102. **(C)** Cell death assays of sgKDM6A/B or sgCtrl/Ctrl HOG EV cells treated with GLIO-1 (*n* = 3). 50 nM: p=0.0053; 100 nM: p=0.0007. **(D)** Doubling times of the indicated bladder cancer cell lines (*n = 3*). **(E)** Cell death assays of endogenous KDM6A-mutant or KDM6A wildtype bladder cancer cell lines treated with ceralasertib for 1–2 population doublings (*n* = 3 per cell line), KDM6A-mutant vs. KDM6A-WT: p=0.0003 **(F)** Immunoblot analysis of sgKDM6A or sgCtrl HTB-5 cells. **(G)** Indel size distribution of amplicons encompassing the target region of the KDM6B sgRNA in HTB-5 cells harboring sgKDM6A/B co-knockout and control sgRNAs. **(H)** Representative photomicrographs of KDM6A/B knockout or control HTB-5 cells treated with 50 nM GLIO-1 or DMSO. Scale bars: 200 µm. **(I)** Immunoblot analysis of sgKDM6A or sgCtrl and **(J)** indel size distribution of sgKDM6A/B or sgCtrl/Ctrl amplicons in HTB-3 cells. **(K)** Representative photomicrographs of KDM6A/B knockout or control HTB-3 cells treated with GLIO-1 or DMSO. Scale bars: 200 µm. **(L)** Immunoblot analysis of sgKDM6A or sgCtrl. **(M)** Indel size distribution of sgKDM6A/B or sgCtrl/Ctrl amplicons in HT1376 cells. **(N–O)** Cell death assays of HT1376 cells harboring KDM6A knockout or KDM6A/B co-knockout and their respective control sgRNAs treated with BAY 2402234 (*n* = 3) (N) or GLIO-1 (O). 10 nM BAY 2402234: sgKDM6A vs. sgCtrl p=0.0002; sgKDM6A/B vs. sgCtrl/Ctrl: p=0.0029. 50 nM GLIO-1: sgKDM6A vs. sgCtrl p=0.039; sgKDM6A/B vs. sgCtrl/Ctrl: p=0.0061. Data are means ± SEM.

**Table S1. sgRNA sequences and corresponding genes comprising 2-oxoglutarate-dependent enzyme sgRNA library, related to Figure 1.**

**Table S2. Results from 2OG-dependent enzyme library CRISPR knockout screen (10 nM BAY 2402234 vs. DMSO) at the indicated time points, related to Figure 1.**

**Table S3. Results from 2OG-dependent enzyme library CRISPR knockout screen (30 µM ceralasertib vs. DMSO), related to Figure 1.**

**Table S4. Results from 2OG-dependent enzyme library CRISPR knockout screen (DMSO vs. library pDNA), related to Figure 1.**

**Table S5. sgRNA sequences comprising base editor sgRNA library, related to Figure 3.**

**Table S6. Results from A>G base editor library (30 µM ceralasertib vs. DMSO), related to Figure 3.**

**Table S7. Results from C>T base editor library (30 µM ceralasertib vs. DMSO), related to Figure 3.**
